# Single-molecule observations of human small heat shock proteins in complex with aggregation-prone client proteins

**DOI:** 10.1101/2024.02.08.579576

**Authors:** Lauren Rice, Nicholas Marzano, Dezerae Cox, Antoine van Oijen, Heath Ecroyd

## Abstract

Small heat shock proteins (sHsps) are molecular chaperones that act to prevent the aberrant aggregation of misfolded proteins. Whilst it is widely suggested that sHsps prevent aggregation by binding to misfolded client proteins, the dynamic and heterogeneous nature of sHsps has hindered attempts to establish the mechanistic details of how sHsp-client protein complexes form. Single-molecule approaches have emerged as a powerful tool to investigate dynamic and heterogeneous interactions such as those that can occur between sHsps and their client proteins. Here, we use total internal reflection fluorescence microscopy to observe and characterise the complexes formed between model aggregation-prone client proteins [firefly luciferase (FLUC), rhodanese, and chloride intracellular channel 1 protein (CLIC)], and the human sHsps αB-crystallin (αB-c; HSPB1) and Hsp27 (HSPB5). We show that small (monomeric or dimeric) forms of both αB-c and Hsp27 bind to misfolded or oligomeric forms of the client proteins at early stages of aggregation, resulting in the formation of soluble sHsp-client complexes. Stoichiometric analysis of these complexes revealed that additional αB-c subunits accumulate onto pre-existing sHsp-client complexes to form larger species - this does not occur to the same extent for Hsp27. Instead, Hsp27-client interactions tend to be more transient than those of αB-c. Elucidating these mechanisms of sHsp function is crucial to our understanding of how these molecular chaperones act to inhibit protein aggregation and maintain cellular proteostasis.

## Introduction

Small heat shock proteins (sHsps) are a ubiquitously expressed, highly conserved class of intracellular molecular chaperones that are characterised by the ability to inhibit aggregation of misfolded proteins in an ATP-independent manner [1,2]. The expression of some sHsps is dramatically upregulated in response to cellular stress, conditions conducive to protein misfolding and aggregation [3,4]. sHsps inhibit this aggregation by binding to, and forming complexes with, aggregation-prone client proteins, trapping them in a state wherein they are not able to form irreversible aggregates [5,6]. As evidence of the critical role of sHsps in maintaining protein homeostasis (proteostasis), dysfunction of the human sHsps HSPB1 (Hsp27) and HSPB5 (αB-c) has been implicated in a number of diseases linked to protein aggregation [7]. Mutations in the *HSPB1* gene encoding Hsp27 are associated with motor neuropathies such as Charcot-Marie-Tooth disease [8] and distal hereditary motor neuropathies [9–11], and overexpression of Hsp27 in an Alzheimer’s disease mouse model reduces amyloid-β plaques and improves spatial learning and memory [12]. Similarly, mutations in the *HSPB5* gene encoding αB-c [13], a major component of the eye lens, result in changes to its’ oligomerisation state and a reduced ability to prevent aggregation [14,15], and are causative of cataract [16].

The oligomeric, at times polydisperse, and dynamic nature of some sHsps is chiefly responsible for the lack of conclusive mechanistic information about how these molecular chaperones prevent protein aggregation [17]. In solution, many sHsps, including αB-c and Hsp27, exist in equilibrium between large molecular mass oligomers (αB-c readily forms ∼18-30mers [17–19], Hsp27 ∼20-28mers [20–22]) and smaller dissociated species (typically monomers and/or dimers). The dynamic equilibrium between sHsp oligomeric forms is impacted by post-translational modifications [23–25] and environmental changes [26–28], which can stimulate disassembly of the oligomers into smaller subunits [29,30]. Furthermore, large sHsp oligomers in complex with a client can be conformationally different to those not associated with a client [31]. This highlights the heterogeneous nature of the sHsps and the complexes they form with client proteins, and the difficulty in interrogating the underlying mechanism by which sHsps interact with aggregation-prone proteins.

Previous work suggests that the mechanism by which sHsps form complexes with aggregation-prone client proteins may be client protein specific. The complexes between sHsps and various clients have been previously investigated using a range of techniques (e.g., *in vitro* light scatter assays [32], co-immunoprecipitation [32] and size-exclusion chromatography [6,26,33]). The stoichiometry of the sHsp-client complexes formed were observed to vary, depending on the type of sHsp tested (e.g., αB-c [32], Hsp27 [32] or related homologs Hsp25 [6], Hsp26 [6], and Hsp18.1 [26,33]), and the aggregation-prone model client protein (e.g., citrate synthase [6,26], rhodanese [6], or firefly luciferase (FLUC) [26]). Furthermore, it has been reported that, of the human sHsps HSPB1-8, αB-c and Hsp27 are the most efficacious and promiscuous when it comes to inhibiting protein aggregation [32]. Moreover, it has been suggested that the molecular mass and stability of the client protein play a key role in the composition of the resulting sHsp-client complexes, and the efficacy of the sHsp against aggregation [26,32,33]. However, these previous studies were limited to studying the complexes that had formed at the end-point of the assay as the techniques used did not allow interrogation of how early-stage sHsp-client protein complexes form and change over time. It remains to be resolved whether there is a shared mechanism by which these sHsps interact with client proteins and how the client protein may affect the composition of these complexes.

Single-molecule fluorescence techniques are particularly amenable to investigating dynamic and heterogeneous biological systems; thus, they are ideally suited for investigating the mechanism by which sHsps interact with client proteins [34,35]. In particular, total internal reflection fluorescence (TIRF)-microscopy facilitates the observation and elucidation of the stoichiometries of heterogeneous protein-protein complexes. We have recently reported the development of a single-molecule fluorescence-based technique that enabled the quantification of stable complexes formed between human αB-c and the model client protein, chloride intracellular channel 1 (CLIC), over time [36]. Here, we further exploit this technique to explore how complexes formed between human sHsps (αB-c and Hsp27) and various client proteins (FLUC, CLIC and rhodanese) evolve over time and whether there are client-dependent differences in the complexes that are formed. In doing so, we show that, once formed, there is an increase in the number of sHsp subunits in a sHsp-client complex over time; significantly more αB-c subunits are recruited into these complexes than Hsp27 subunits. Furthermore, we demonstrate that there are client-specific differences in the complexes formed between these sHsps and aggregation-prone proteins, potentially due to differences in the molecular mass and/or hydrophobicity of the clients. Our results provide evidence for a common underlying mechanism by which sHsps form complexes with client proteins, whilst highlighting the heterogeneity in the sHsp-client protein complexes that are formed.

## Results

### The human sHsps αB-c and Hsp27 inhibit the aggregation of destabilised client proteins in vitro

Traditionally, ensemble-averaging measurements have been used to monitor the aggregation of client proteins and the ability of sHsps to inhibit this aggregation [6,37,38]. As such, we validated our investigation into the chaperone function of αB-c and Hsp27 against CLIC, FLUC and rhodanese by measuring the change in light-scatter over time for each client protein (indicative of amorphous aggregate formation) in the presence and absence of the sHsps.

When each client was incubated in the absence of sHsps an increase in light scatter was observed over time, indicating that aggregation had occurred (Fig. 1A-C). Notably, when the clients were incubated in the presence of αB-c, there was a significant decrease in light scatter observed for CLIC and FLUC (p < 0.005), but not rhodanese, when compared to the control protein, ovalbumin (Fig. 1D). In contrast, when the clients were incubated in the presence of Hsp27, there was a significant reduction in light scatter for all three client proteins, consistent with the inhibition of protein aggregation (Fig. 1E). This data is in agreement with previous reports that αB-c and Hsp27 do not inhibit the aggregation of all clients comparably [6,32], which could be due to mechanistic differences by which these sHsps form interactions with their clients. Such interactions might be influenced by differences in the oligomerisation state and rate of subunit exchange between these two human sHsps [22,39], as well as the exposed hydrophobicity of the client during misfolding and/or the rate and propensity of client aggregation (Fig. 1A-C). Whilst the presence of ovalbumin did lead to a decrease in light scatter for all three clients tested, this did not occur to the same extent as for the sHsps, and is consistent with previous studies that have shown that albumin demonstrates weak chaperone-like activity against some clients [40,41].

**Figure 1.**
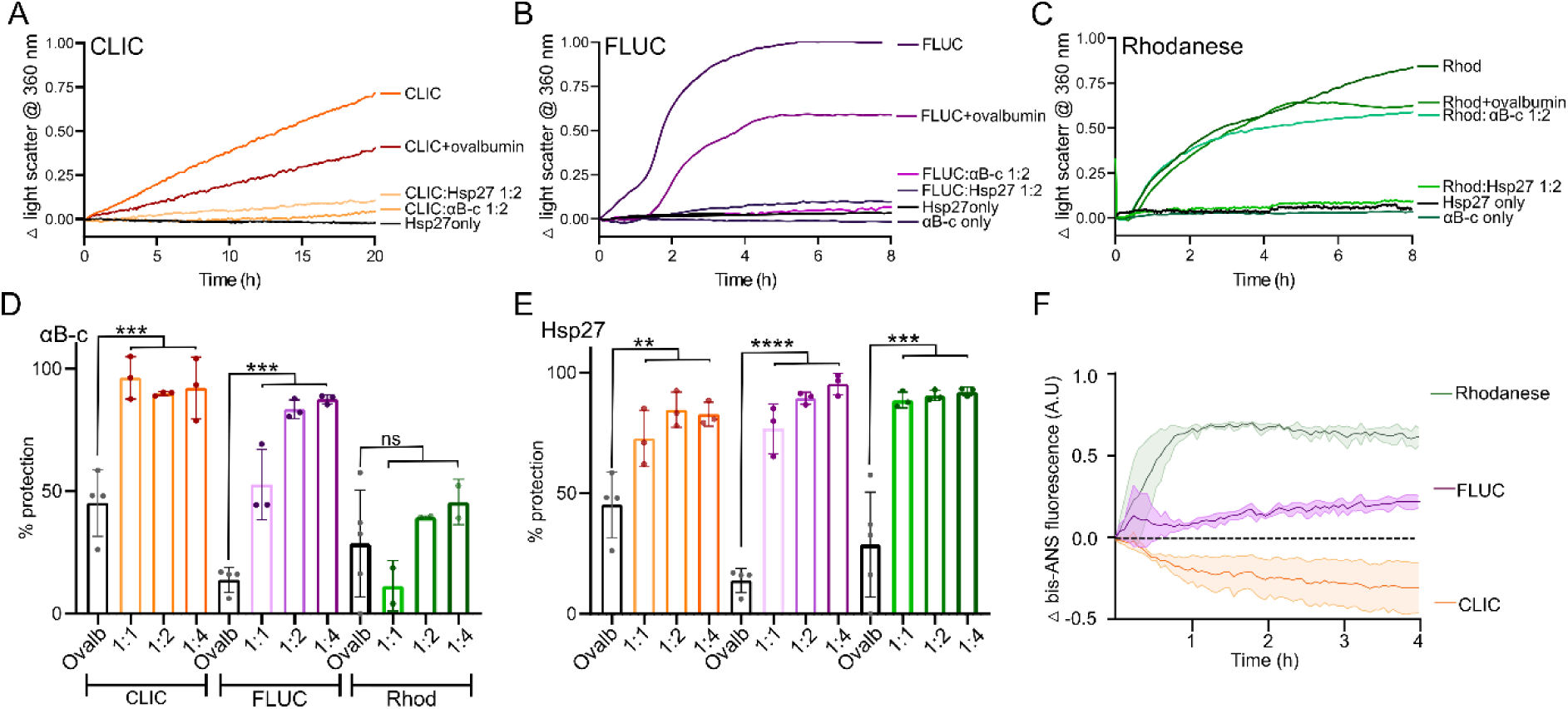
Differences in the capacity of the human sHsps, αB-c and Hsp27, to inhibit the heat-induced amorphous aggregation of client proteins. (A-C) Example light scatter traces from the heat-induced aggregation of the client proteins. Recombinant client protein (A-CLIC at 30 µM, B-FLUC at 4 µM, or C-rhodanese at 4 µM) was incubated at 42°C for 20 h (CLIC), or 8 h (FLUC and rhodanese) in the presence or absence of a 1:2 molar ratio (client:sHsp) of either αB-c or Hsp27, or the control protein ovalbumin, and the change in light scatter at 360 nm over time monitored. **(D-E)** The percent reduction in aggregation (light scatter) of CLIC, FLUC and rhodanese afforded by varying molar ratios of αB-c (D) and Hsp27 (E), and the control protein ovalbumin at the highest molar ratio tested (1:4, client:ovalbumin). Data reported is the average ± S.D. of three independent experiments. Data were analysed by one-way ANOVA and Tukey’s post-hoc test, where ****, ***, and ** indicates p < 0.0001, 0.001, and 0.01, respectively; ns indicates p > 0.05. **(F)** Exposed hydrophobicity of client proteins was monitored by bis-ANS fluorescence. CLIC, FLUC and rhodanese (200 nM) were incubated at 42°C for 6 h in the presence of 20 µM bis-ANS; fluorescence emission from bis-ANS at 480 nm was monitored over time and reported as the average change ± S.E. of three independent experiments.

We next investigated whether the interaction of sHsps with these model client proteins correlates with the amount of exposed hydrophobicity on the client during heat-induced unfolding; to do so, we performed 1,1′-bi(4-anilino)naphthalene-5,5′-disulfonic acid (bis-ANS) assays (Fig. 1F). When FLUC and rhodanese were incubated with bis-ANS under the same conditions used for the aggregation assays (Fig. 1A-C), there was an increase in bis-ANS fluorescence over time that is indicative of more surface-exposed hydrophobicity upon client misfolding. However, the increase in fluorescence occurred faster and to a greater extent for rhodanese compared to FLUC. In contrast, bis-ANS fluorescence decreased upon incubation with CLIC, which suggests that CLIC misfolding results in hydrophobic regions becoming more buried when compared to the native state.

Collectively, these data confirm that sHsps inhibit the aggregation of these client proteins, and that the extent to which this occurs varies depending on the client. Differences in the capacity of αB-c and Hsp27 to inhibit the aggregation of the various clients could relate to varying degrees of surface-exposed hydrophobicity during the early stages of client misfolding. However, specific mechanistic differences that may underlie these observations were not able to be determined using these techniques; as such, we turned to a single-molecule approach to further interrogate these sHsp-client protein interactions.

### The number of αB-c subunits in complex with client proteins increases over time and is client-dependent

To interrogate the interaction between the sHsps αB-c and Hsp27 with these model client proteins, we utilised a previously described single-molecule photobleaching approach that enables the identification and characterisation of stable sHsp-client complexes [36]. Briefly, this approach entails Alexa Fluor 647 (AF647)-labelled client proteins being incubated in heat-denaturing conditions in the absence or presence of Alexa Fluor 488 (AF488)-labelled sHsps. Aliquots are taken over the course of the reaction (Fig. 2A), crosslinked to prevent complex dissociation (Supp Fig. 1), diluted and immobilised on a coverslip for imaging using TIRF microscopy until all fluorophores are photobleached. The spectrally distinct fluorophores attached to the client or sHsp are visible in separate imaging channels which, when aligned and overlaid, enable identification of coincident fluorescent spots indicative of sHsp-client complexes (Fig. 2A).

**Figure 2.**
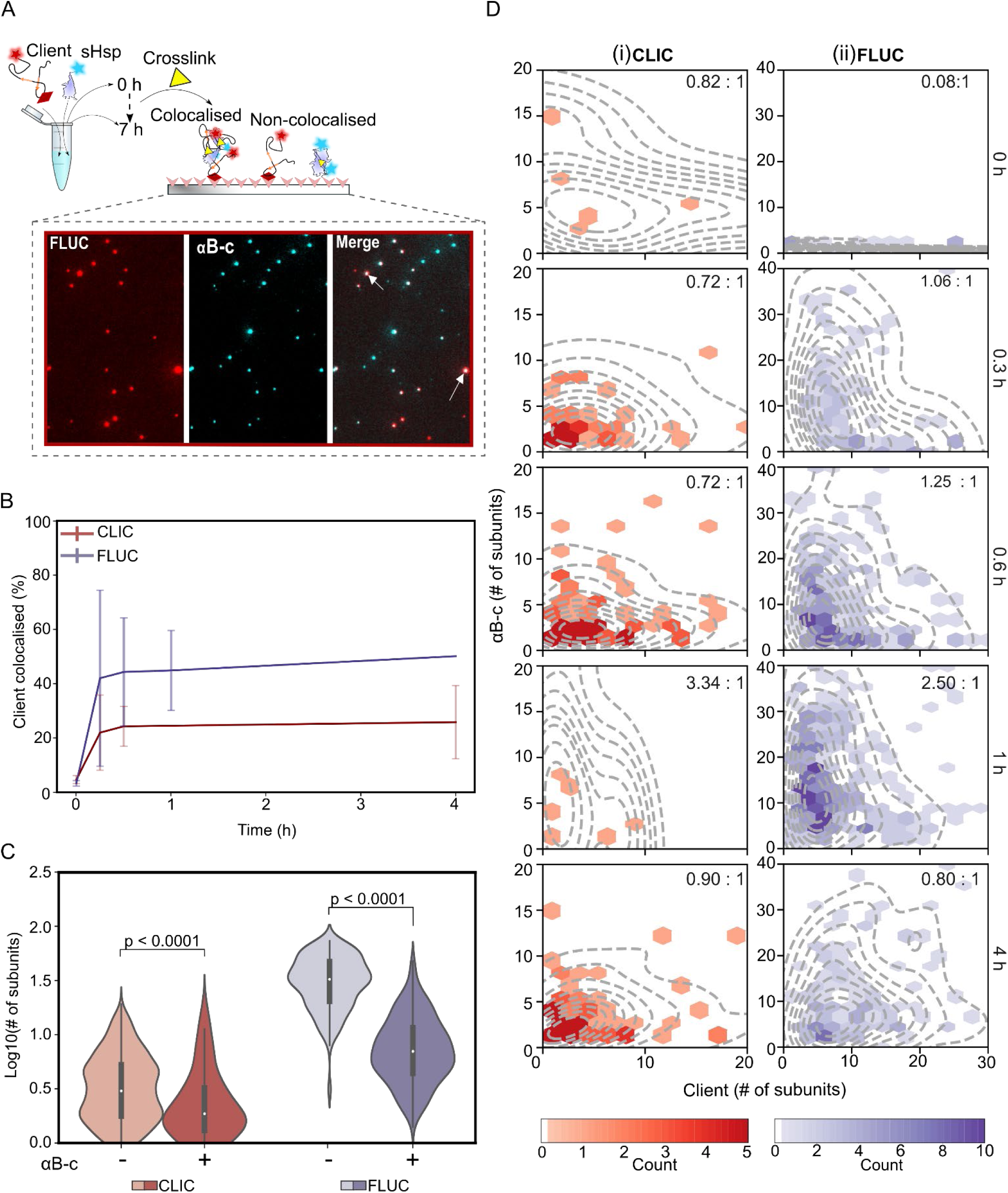
The number of αB-c subunits within sHsp-client protein complexes increases over time in a client-dependent manner. αB-c-AF488 and CLIC-or FLUC-AF647 were incubated (42°C for up to 7 h) in the absence or presence of αB-c (2:1 molar ratio; αB-c:client); aliquots were taken at time points throughout the incubation. Following incubation, samples were immediately crosslinked, diluted, and incubated in flow cells for 15 min before imaging using TIRF microscopy. **(A)** Schematic depicting the workflow used to identify sHsp:client complexes formed during incubation. Representative TIRF microscopy image shows the fluorescence emission from FLUC-AF647 and αB-c-AF488 when incubated together, observed in separate emission channels. Overlaying these channels (‘Merge’) allows identification of colocalised (white arrows) and non-colocalised molecules. **(B)** The percentage (%) of CLIC-AF647 (red) or FLUC-AF647 (purple) molecules colocalised with αB-c-AF488 at each time point, reported as the mean ± S.D. of 3 independent replicates. **(C)** Violin plots show the distributions of CLIC-AF647 (red) and FLUC-AF647 (purple) molecule sizes (log_10_ number of subunits/molecule) after 7 h incubation with (‘+’) or without (‘-’) αB-c-AF488. Molecule size reported includes all molecules (i.e. both colocalised and non-colocalised with αB-c). Results include measurements from 3 independent experiments and, where marked, statistical comparisons between distributions was performed via Kruskal-Wallis test for multiple comparisons with Dunn’s procedure (p values indicated). **(D)** Hexbin plots show the relative abundance of protein subunits and median molar ratio (sHsp:client) (inset) of αB-c-AF488 (y-axis) and CLIC-AF647 (i) or FLUC-AF647 (ii) (x-axis) within complexes at each time point during incubation. Each hexbin plot is overlaid with the kernel probability density (dashed line) of complexes at each time point. Data shown is from all molecules in complexes measured across the 3 independent experiments.

We used this approach to investigate the sHsp-client combinations where aggregation was significantly inhibited i.e., αB-c with either FLUC or CLIC; Hsp27 with all three clients (Fig. 1). Within 0.3 hours, ∼20% of immobilised CLIC molecules were colocalised with αB-c and this proportion of colocalisation remained static throughout the remainder of the time course (Fig. 2B). This is consistent with our previous observations without crosslinking [36], indicating that the chemical crosslinker does not alter complex formation. Similarly, 40% of FLUC molecules were colocalised with αB-c within 0.3 hours (Fig. 2B). The increased formation of colocalised complexes between αB-c and FLUC compared to CLIC is consistent with sHsps preferentially interacting with clients with increased hydrophobicity and aggregation propensity (Fig. 1F). However, despite a two-fold molar excess of chaperone, 100% colocalisation was not achieved, suggesting sHsps are selective for a sub-population of client molecules. This sub-population may be those that misfold first and/or are more aggregation-prone.

Using this single-molecule approach it was also possible to determine the size of client species when incubated in the presence or absence of αB-c. When CLIC or FLUC were incubated with αB-c, the client species observed were significantly smaller than those incubated in the absence of αB-c (p < 0.0001). Given that complexes form in this time (Fig. 2B), this indicates that the formation of complexes acts to inhibit aggregation of both clients (Fig. 2C). As expected, the FLUC species were smallest prior to incubation indicating that any increase in size is indeed due to aggregation (Supp Fig. 2). Interestingly, the distribution of client molecule sizes when incubated with αB-c was also broader than those seen prior to incubation (Supp Fig. 2). This indicated that some molecules were still able to increase in size despite a proportion of client molecules forming complexes with αB-c.

We next determined the composition of the sHsp-client complexes by calculating the ratio of client to sHsp molecules within individual complexes over time. When αB-c was in complex with CLIC, the relative molar ratio of αB-c:CLIC fluctuated from 0.72:1 to 3.34:1 over time (Fig. 2D-i), although this change was not statistically significant (p = 0.24). However, very few αB-c:CLIC complexes were observed at each time point. This suggests that, given the surface-exposed hydrophobicity of CLIC decreases under these conditions (Fig. 1F), formation of complexes between αB-c and CLIC is not favoured. The distribution of αB-c:FLUC molar ratios within complexes that did form increased significantly over time (p < 0.0001), and the median increased from 0.08:1 (prior to incubation i.e., 0 hours) to 2.5:1 after 1 hour (Fig. 2D-ii). Crucially, FLUC species in complex with αB-c remained significantly smaller than FLUC species incubated alone (Fig. 2C), indicating that the formation of complexes and the subsequent recruitment of additional αB-c subunits into the complex acts to prevent large FLUC aggregates from forming (Fig. 2D). Interestingly, substantially fewer αB-c-FLUC complexes were present after 4 hours of incubation, and the population of complexes that contained a large number of αB-c subunits (more than 20) were not observed. This resulted in the median molar ratio of αB-c-FLUC complexes decreasing 3-fold from 2.5:1 (αB-c:FLUC, at 1 hour) to 0.8:1 (at 4 hours).

These data indicate that αB-c initially forms stable complexes with FLUC and then additional αB-c subunits are recruited into these complexes over time; in doing so, extensive aggregation of FLUC is inhibited. The subsequent decrease in the total number and relative molar ratio of αB-c-FLUC complexes at later time points suggests that, within the largest complexes, the manner by which the αB-c subunits associate with the FLUC molecules prevents them from binding the coverslip. Whilst it could be argued that the decrease in molar ratio within complexes is due to αB-c-FLUC complexes dissociating over time, this would not result in the observed decrease in the total number of FLUC molecules bound to the coverslip. As such, only those FLUC molecules associated with fewer αB-c subunits are able to be detected at this time point (4 hours) and no complexes were detected at later times.

Taken together, this data shows that αB-c forms stable complexes with both CLIC and FLUC. Differences in the extent to which complexes are formed are likely due to differences in hydrophobicity between the two clients. To determine whether these findings are unique to αB-c or are a more general mechanism by which human sHsps interact with client proteins, similar experiments were performed with Hsp27.

### Hsp27 prevents client aggregation through complex formation and transient interactions

We next sought to determine whether the interaction between Hsp27 and the model client proteins CLIC, FLUC and rhodanese was the same as for αB-c. As for αB-c, the percentage of CLIC molecules that colocalised with Hsp27 increased in the first 1 hour to 15% (Fig. 3A). Notably, Hsp27 more rapidly colocalised with rhodanese when compared to CLIC, and 40% of rhodanese molecules were colocalised within 0.3 hours (Fig. 3A). Colocalisation of Hsp27 with both rhodanese and CLIC plateaued after 1 hour (Fig. 3A). When Hsp27 was incubated with FLUC, the proportion of complexes was dynamic; within the first 0.6 hours of incubation there was a rapid increase in colocalisation to ∼35%. This was followed by a two-fold decrease in colocalisation (to ∼17%) at 1 hour. Unlike αB-c (Fig. 2), there was no decrease in the number of observed FLUC molecules bound to the coverslip surface. The colocalisation of FLUC with Hsp27 plateaued after this point. The unexpected observation of rapid Hsp27 binding and apparent complex dissociation with FLUC suggests a more dynamic interaction compared to those between FLUC and αB-c. Collectively, these results show that despite Hsp27 inhibiting the aggregation of all clients (Fig.1), colocalisation with client molecules varies over time and between clients. As such, there may be subtle differences in the mechanism by which Hsp27 forms complexes with different clients.

**Figure 3.**
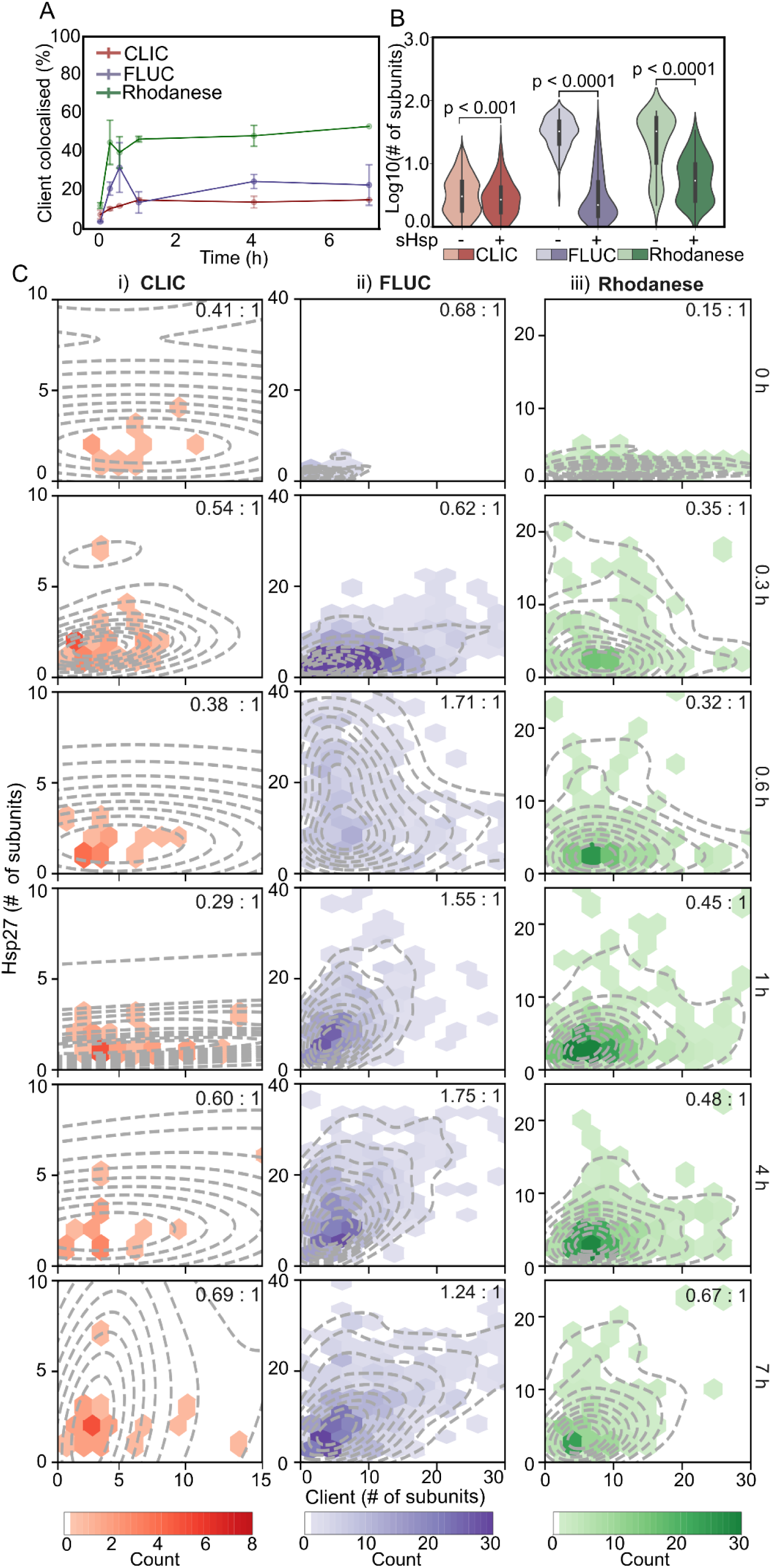
The number of subunits of Hsp27 within complexes increases in a client-dependent manner. Hsp27-AF488 and the client proteins (CLIC-, FLUC-or rhodanese-AF647) were incubated (42°C for up to 7 h) in the absence or presence of Hsp27 (2:1 molar ratio, Hsp27:client); aliquots were taken at time points throughout the incubation. Following incubation, samples were immediately crosslinked, diluted, and incubated in flow cells for 15 min before imaging using TIRF microscopy. **(A)** The percentage (%) of CLIC-AF647 (red), FLUC-AF647 (purple) or rhodanese-AF647 (green) molecules colocalised with Hsp27-AF488 at each time point, reported as the mean ± S.D. of 3 independent replicates. **(B)** Violin plots show the distributions of CLIC-AF647 (red), FLUC-AF647 (purple), and rhodanese-AF647 (green) molecule sizes (log_10_ number of subunits/molecule) after 7 h incubation with (‘+’) or without (‘-’) Hsp27-AF488. Molecule size reported includes all molecules (i.e. both colocalised and non-colocalised with Hsp27). Results include measurements from 3 independent experiments and, where marked, statistical comparisons between distributions was performed via Kruskal-Wallis test for multiple comparisons with Dunn’s procedure (p values indicated). **(C)** Hexbin plots show the relative abundance of protein subunits and median molar ratio (sHsp:client) (inset) of Hsp27-AF488 (y-axis) and each of the clients (CLIC-AF647 (i), FLUC-AF647 (ii) and rhodanese-AF647 (iii)) (x-axis) within complexes at each time point during incubation. Each hexbin plot is overlaid with the kernel probability density (dashed line) of complexes at each time point. Data shown is from all molecules in complexes measured across the 3 independent experiments.

To interrogate this, we first determined the effect of Hsp27 on the molecule size of each client species regardless of whether it was colocalised with Hsp27 or not (Fig. 3B). Consistent with results observed for αB-c, the size distribution of all three clients was significantly smaller when incubated in the presence of Hsp27 compared to when Hsp27 was not present, indicating that Hsp27 inhibited the aggregation of these proteins (p<0.0001). Again, as expected, the FLUC and rhodanese molecules were small prior to incubation (Supp Fig. 2), and thus the increase in size in the absence of Hsp27 was attributable to aggregation (Fig. 3). Additionally, the distribution of CLIC and FLUC molecule sizes when incubated with Hsp27 was broader than those seen prior to incubation (Supp Fig. 3). Taken together with the finding that Hsp27 only forms complexes with a sub-population of the client molecules (Fig. 3A), we sought to better understand which population of client molecules is sequestered into complexes with Hsp27. As such, we examined the composition of and changes within the Hsp27:client complexes over time.

The stoichiometric distribution of complexes between CLIC and Hsp27 ranged from a median molar Hsp27:CLIC ratio of 0.29:1 – 0.69:1 (Fig. 3C-i). However, as with αB-c and CLIC, there were very few stable Hsp27-CLIC complexes observed in this sample and, as such, changes within a single complex influenced the median substantially. Notably, the number of FLUC or rhodanese molecules that were in complexes with Hsp27 did not change substantially over the incubation, indicating that formation of these complexes inhibits the extensive aggregation of the client. However, there were substantial differences in the molar ratio of the Hsp27:client complexes that formed over time between rhodanese and FLUC (Fig. 3E-ii-iii). Whilst only smaller species of Hsp27 (i.e., monomers or dimers) were present in complex with FLUC at early time points, the number of Hsp27 molecules in complexes increased ∼ 2.5 fold within 4 hours (from 0.68:1 to 1.75:1, p < 0.0001). However, the molar ratio of Hsp27:FLUC complexes significantly decreased by the final (7 hour) time point to 1.24:1 (p < 0.05), indicating that the recruitment of Hsp27 subunits into these complexes does not continue to occur after a point. Importantly, even at the maximum average molar ratio of Hsp27:FLUC, the number of Hsp27 subunits within complexes was still only half that of αB-c under the same conditions (Fig. 2). Prior to incubation (0 hours) monomers and dimers of Hsp27 bound to a distribution of rhodanese oligomers (1 – 20 subunits, median molar ratio of 0.15:1). The molar ratio of Hsp27:rhodanese complexes increased steadily throughout the incubation, reaching a maximum of 0.67:1 after 7 hours, indicating recruitment of Hsp27 molecules into complexes with rhodanese. However these complexes always contained less Hsp27 subunits per rhodanese molecule compared to those complexes formed with FLUC.

In summary, these data show that, similar to αB-c, Hsp27 binds CLIC, FLUC and rhodanese over time to different extents, supporting our proposition that differences in hydrophobicity between the clients may govern the extent of complex formation. In contrast to αB-c, Hsp27 displayed dynamic complex formation with FLUC, and formed proportionately fewer complexes with less Hsp27 molecules per FLUC monomer. Interestingly, incubation of FLUC with αB-c resulted in the disappearance of large complexes after 1 hour, and most molecules overall by 4 hours, suggesting this complex formation eventually prevents FLUC binding to the coverslip. In contrast, Hsp27:FLUC complexes and non-colocalised FLUC molecules were abundant and small for the entire 7 hour incubation, suggesting that complex formation with Hsp27 does not prevent coverslip binding. It is clear that both sHsps inhibit the formation of large aggregates of FLUC, and that Hsp27 also prevents the formation of large aggregates of rhodanese. The number of αB-c and Hsp27 subunits in complexes increases over time, but the number of client molecules in the complexes remain relatively stable, leading to an increase in the sHsp:client molar ratio of the complexes.

### Hsp27 and αB-c maintain client proteins in small oligomeric states

To better understand how the sHsps recognise and maintain the entire population of client protein molecules in a non-aggregated state (Fig. 2B, 3A), we directly compared the size of the non-colocalised client molecules at each time point with those in complexes (Fig. 4A and C). The number of CLIC subunits in complex with αB-c was similar throughout the incubation period to those not in complex (p > 0.05) (Fig. 4C-i). In contrast, at early time points (i.e. <1 h) the number of FLUC subunits per molecule was significantly higher when in complex with αB-c compared to those not in complex (p < 0.01) (Fig. 4C-ii), suggesting that αB-c selectively binds small oligomeric species rather than native or misfolded monomers. The size of the non-colocalised FLUC molecules increased significantly to ∼ 10 subunits after 4 hours (p < 0.01, Supp Fig 4A-ii); in contrast, the colocalised FLUC molecules were similar in size throughout the incubation (∼ 8 subunits, p > 0.05, Supp Fig4A-ii). Similarly, for CLIC, only the non-colocalised molecules increased significantly during the first hour of incubation, before decreasing to the same size as those present prior to incubation (Supp Fig. 4A-i, p < 0.05). This suggests that the initial associations between CLIC molecules are not stable. Together, this data indicates that client proteins that are not bound by a sHsp grow in size.

**Figure 4.**
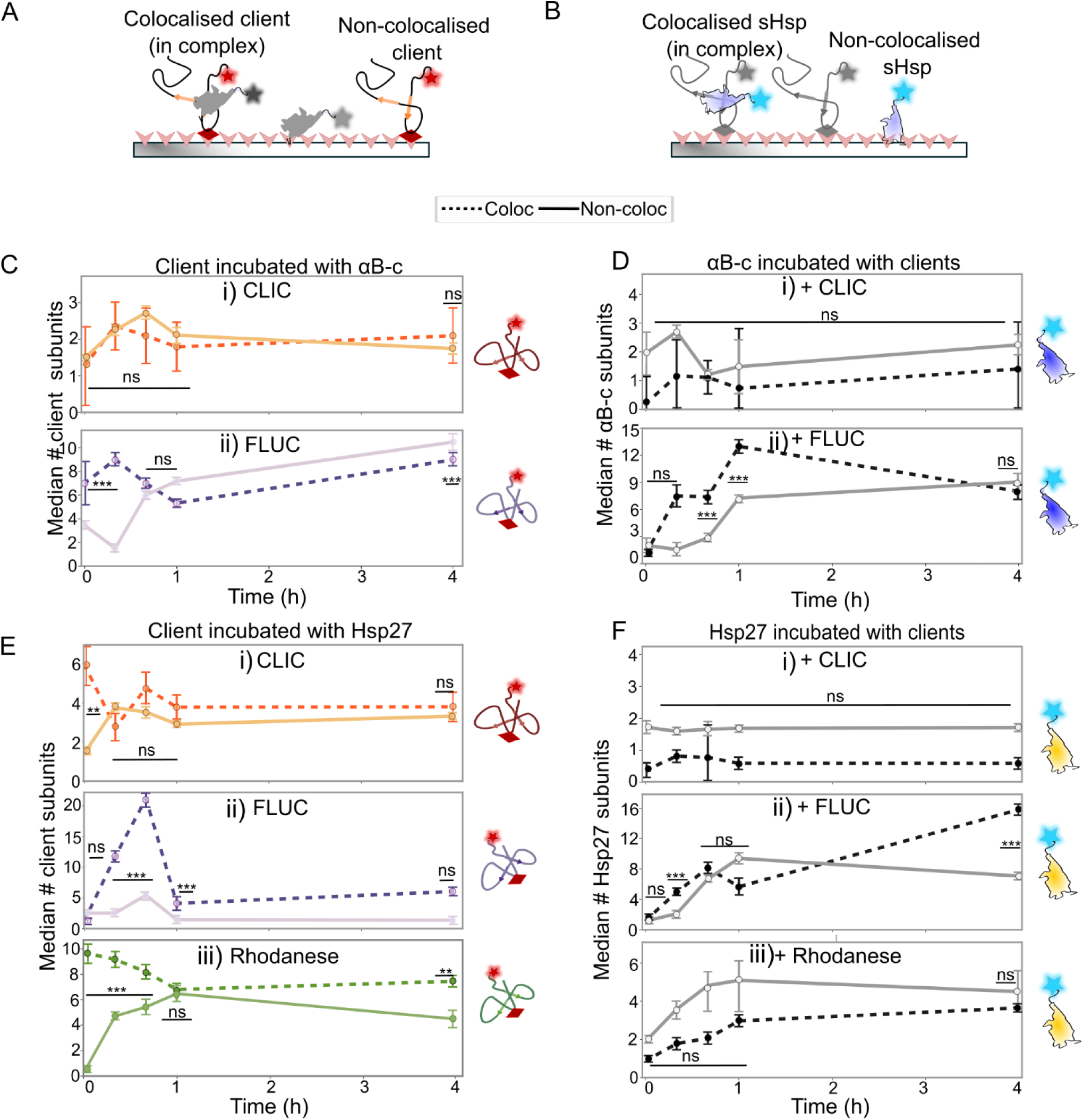
sHsps maintain client proteins in small oligomeric states. AF488-labelled sHsps (αB-c or Hsp27) were incubated with the AF647-labelled client proteins (CLIC, FLUC or rhodanese) (42°C for 4 h, 2:1 molar ratio) and aliquots were taken throughout the incubation. Samples were immediately crosslinked, diluted, and incubated in flow cells for 15 min before imaging using TIRF microscopy. The molecule size (# of subunits/molecule) was calculated for all molecules, and the data was filtered for the **(A)** client and **(B)** sHsp molecules that were colocalised with one another (i.e., in complexes) and those that were not colocalised (i.e., not in complexes). (**C - E**) The median (± S.E.) number of subunits per molecule of both colocalised (‘Coloc’) and non-colocalised (‘Non-coloc’) molecules was plotted at each time point for the clients CLIC (i), FLUC (ii) or rhodanese (E-iii) incubated with (C) αB-c or (E) Hsp27. The size of the sHsps in these treatments are shown in the corresponding plots for (D) αB-c and (F) Hsp27. The median is calculated at each time point from all molecules from 3 independent experiments. Comparisons were made via a two-way ANOVA with Tukey’s HSD post-hoc testing performed to compare the number of subunits between colocalisation states (i.e. ‘Coloc’ versus ‘Non-coloc’) at each time point; ns = not significant (p > 0.05); *** = p < 0.0005; ** = p < 0.005.

When we compared the number of αB-c subunits per molecule when incubated with CLIC (Fig. 4D-i), we observed that small αB-c oligomers initially bind to CLIC. These do not increase over time (Supp Fig. 4B-i, p > 0.05), and are similar between those that are colocalised and not-colocalised (Fig. 4D-i, p > 0.05). When incubated with FLUC, the number of colocalised αB-c subunits per complex increased rapidly in the first hour (∼13 subunits/complex after 1 hour) (Fig. 4D-ii, Supp. Fig 4B-ii, p < 0.05). Interestingly, the size of non-colocalised αB-c also increased between 0.6 – 4 hours, however this occurred at a slower rate and to a lesser extent (∼ 6 subunits/oligomer after 1 hour) compared to the colocalised αB-c (Supp Fig. 4B-ii, p < 0.05). By the end of the incubation, the colocalised and non-colocalised αB-c molecules were approximately the same size (Fig. 4D-ii, p > 0.05). These data suggest that αB-c binds to FLUC molecules that have already begun to aggregate; over time, small αB-c subunits are recruited into complexes such that, at later time points, there are fewer subunits available to inhibit the further aggregation of FLUC. Since this work has demonstrated that the smaller chaperone species (i.e., monomers/dimers) preferentially associate with the misfolded protein, it is likely that these sHsp species are depleted from solution first and sequestered into sHsp:client complexes. This would account for the observed increase in median size of non-colocalised αB-c (Fig. 4D-ii).

Next, we compared the size of CLIC, FLUC and rhodanese molecules that had been incubated with Hsp27 (Fig. 4Ei-iii). The number of CLIC (Fig. 4E-i) or rhodanese (Fig. 4E-iii) subunits was initially higher when in complex with Hsp27 compared to those not in complex with Hsp27 (0 – 0.6 hours, p < 0.05). However, the median number of rhodanese molecules within these complexes decreased at later time points (> 1 hour, Supp Fig. 4C-iii, p < 0.05), whilst the number of CLIC molecules within complexes with Hsp27 did not change (Supp Fig. 4C-i; p > 0.05). Conversely, the median size of the client molecules (both CLIC and rhodanese) not in complex with Hsp27 increased over time (Supp Fig. 4C-i, iii) such that they were approximately the same size as those molecules in complex (Fig.4E-i, iii; p > 0.05). This supports the proposal that for these clients, molecules which are not initially bound by a sHsp eventually begin to oligomerise and aggregate.

FLUC predominantly existed as small species (i.e., 2 subunits) when non-colocalised or in complex with Hsp27 prior to incubation (0 hours; Fig. 4E-ii; p > 0.05). However, the median number of FLUC subunits in complexes with Hsp27 increased significantly at each time point until 0.6 hours, reaching up to 20 subunits per molecule (Supp Fig. 4C-ii, p < 0.001). By 1 hour, the number of FLUC subunits in complexes decreased significantly compared to the prior time points (Supp Fig. 4C-ii, p < 0.05), becoming similar in size to that observed for non-colocalised FLUC molecules (∼ 5 FLUC subunits/complex) (Fig 4E-ii). The FLUC molecules that were not in complexes also increased in the first 0.6 hours of incubation (Supp Fig. 4C-ii; p < 0.05), before decreasing in size at each remaining time point (Supp Fig. 4C-ii, p < 0.05). However, the magnitude of these changes was smaller than those in complex with Hsp27. Thus, the non-colocalised molecules remained small (< 5 subunits per FLUC molecule) and contained fewer subunits than those in complex throughout the incubation (Fig. E-iii, p < 0.05). This is in stark contrast to results observed in the presence of αB-c, whereby the non-colocalised FLUC increased significantly in size. Taken together, these data suggest that Hsp27 inhibits aggregation through both stable and transient interactions with aggregation-prone forms of client proteins.

Finally, we compared the number of Hsp27 subunits per molecule when non-colocalised or in complex with clients. Prior to incubation, Hsp27 molecules in complex and not in complex with all clients were monomers and dimers (Fig. 4F-i-iii; p > 0.05). When Hsp27 was incubated in the presence of CLIC (the least aggregation-prone client with the lowest propensity to form complexes), the number of Hsp27 subunits in non-colocalised and colocalised molecules did not change over time (1-2 subunits/molecule) (Supp Fig. 4D-i; p > 0.05). In contrast, when incubated in the presence of FLUC or rhodanese, the number of Hsp27 subunits increased throughout the incubation period (Supp Fig. 4D-ii,iii, p < 0.05) for Hsp27 molecules in complex and not colocalised with the client. For rhodanese, this increase in the number of Hsp27 molecules was more gradual for those in complexes, and resulted in there being no difference between the number of Hsp27 subunits in and out of complexes (Fig. 4F-iii; p > 0.05). This trend was similar for Hsp27 and FLUC, except for at 4 hours, at which point there were more Hsp27 subunits in complex with FLUC than there were within non-colocalised Hsp27 molecules (Fig. 4F-ii; p < 0.05). Similar to αB-c, this could be due to the smaller monomeric and/or dimeric species of Hsp27 being depleted from solution as they bind the client.

These data demonstrate the preferential binding of small oligomers of sHsps to the largest client molecules and the subsequent increase in size of those client molecules not in complexes. Thus, this work indicates that both αB-c and Hsp27 discriminate between aggregated and non-aggregated forms of client proteins and prioritise binding to the aggregated species. The differences in size between the non-colocalised and colocalised molecules during incubation is dependent on the client present, likely due to the difference in the number of sHsp subunits required to inhibit its aggregation (i.e. more for FLUC than CLIC or rhodanese).

## Discussion

It is well known that the sHsps αB-c and Hsp27 form a range of oligomers in the absence of a client, and that when a client is present they can form large, polydisperse sHsp:client complexes [2,26,32,42]. However, it remains to be resolved whether the mechanism by which these sHsps forms complexes with clients is the same, and whether the formation of complexes is influenced by the client protein. Collectively, the results reported in this study provide evidence for a common underlying mechanism by which sHsps form complexes with aggregation-prone client proteins and highlight the heterogeneity of the sHsp-client protein complexes that are formed.

### αB-c and Hsp27 share a common underlying mechanism by which they interact with client proteins to inhibit their aggregation

We propose that, at early time points, αB-c and Hsp27 behave in a similar manner to inhibit aggregation of client proteins; specifically, when sHsps are exposed to increased temperatures, the dynamic exchange of subunits between oligomers increases. Simultaneously, heat induces the client protein to misfold, which results in exposure of hydrophobic regions on the polypeptide, and the formation of small oligomeric/early aggregated species. As such, the affinity of an individual sHsp subunit for the hydrophobic client becomes significantly higher than the affinity for other sHsp subunits. This results in a decrease in size of the larger sHsp oligomers [43,44], increased availability of free sHsp monomers and dimers in solution and drives the formation of sHsp-client complexes. This is consistent with our observations that complex formation for both αB-c and Hsp27 occurs within the first 0.6-1 hours following incubation with a client (Fig 2B-3A). Moreover, complex formation and the extent to which this occurs is enhanced for clients for which there is a significant increase in hydrophobicity during this time (i.e. rhodanese, Fig. 1F) and, as such, may require stable complex formation to inhibit aggregation [45,46]. This is also apparent when comparing the early stage complex formation of Hsp27 with FLUC and rhodanese; Hsp27 forms a greater proportion of stable complexes with rhodanese than it does FLUC, which correlates with the differences in hydrophobicity between these two clients at this time (i.e. rhodanese hydrophobicity increased to a level ten times higher than that of FLUC in the first 1 hour).

The results from this work indicate that, over the initial periods during which sHsp-client complexes form (i.e. up to 1 hour), additional sHsp subunits are recruited from solution into the complexes, resulting in larger complexes with more sHsps bound. The ‘seeding’ or recruitment of these sHsp subunits into complexes has been suggested previously as a mechanism by which αB-c inhibits aggregation [36]. Here, we demonstrate that this is a common underlying mechanism shared by both αB-c and Hsp27. The increase in the number of sHsp subunits in complexes over time is particularly apparent for those client proteins which form larger aggregates in the absence of a sHsp (i.e. FLUC and rhodanese). We propose that the extent to which this recruitment occurs may relate to the molecular mass of the client protein, since the availability of binding sites on a larger client may be greater. Evidence to support this from this work is that the complexes both αB-c and Hsp27 formed with FLUC (the largest client protein, at 62 kDa) had the largest number of sHsp subunits within the complex when compared to complexes formed with CLIC (27 kDa) or rhodanese (35 kDa). Of note, there were consistently more αB-c subunits recruited into αB-c:client complexes than Hsp27 in Hp27:client complexes, despite Hsp27 inhibiting aggregation to a greater or similar extent compared to αB-c; Hsp27 was effective at inhibiting rhodanese aggregation whereas αB-c was not.

### Differences in the complexes formed by αB-c and Hsp27 with client proteins

Differences in the complexes formed between αB-c and Hsp27 with FLUC became most apparent at later time points during incubation (from 1 hour onwards) (complexes formed with CLIC were also interrogated, however, far fewer complexes were formed). When FLUC was incubated with αB-c for greater than 1 hour, fewer complexes were detectable with our single-molecule approach. We surmise that the complexes that were no longer present after 4 hours were those comprised of many αB-c subunits as the distribution of the complexes that were observed after 1 hour shifted to those with fewer αB-c subunits. This suggests that the manner by which αB-c interacts with the FLUC in these complexes may shield regions of FLUC such that it can no longer bind to the coverslip surface [47]. In contrast, the relative proportion of FLUC molecules within Hsp27:FLUC complexes was found to decrease after 1 hour of incubation, suggesting that some of the Hsp27:FLUC complexes dissociate, resulting in more non-colocalised FLUC subunits. Of note, despite evidence for some Hsp27:FLUC complexes dissociating, many large Hsp27:FLUC complexes remained at the end of the incubation (in contrast to αB-c) suggesting that the structural interactions between Hsp27 and FLUC in these complexes are dynamic and heterogeneous.

We were able to show via our single-molecule technique that whilst aggregation (as detected by light scatter) is inhibited overall in the presence of the sHsps, there is a proportion of client molecules not in complex with sHsps that increase in size during incubation. Thus, when FLUC and αB-c, or rhodanese and Hsp27, were incubated together, the non-colocalised molecules increased in size over time, suggesting incomplete inhibition of aggregation. However, when FLUC was incubated with Hsp27, those molecules not in complex remained small throughout the entire incubation. Taken together, these data suggest that the formation of stable complexes with clients is not the sole mechanism responsible for sHsp inhibition of aggregation; transient interactions, such as those that take place between Hsp27 and FLUC, also take place between sHsps and client proteins and these too act to inhibit the client from forming aggregates. Moreover, recruitment of sHsp subunits into complexes (as occurs with FLUC and αB-c) may actually deplete free sHsp subunits from solution such that they are not available to participate in transient interactions, resulting in non-complexed molecules aggregating over time. It has been previously suggested that the interaction of Hsp27 with FLUC may act to facilitate active refolding by the Hsp70 chaperone system [48]. The ability of Hsp27 to readily dissociate from complexes may assist in this process. Additionally, compared to αB-c, Hsp27 subunits have weaker intermolecular affinities, evidenced by a higher propensity for Hsp27 oligomers to dissociate into smaller species upon dilution [2,7,20,30,49] and an increased rate of subunit exchange between homo-oligomers in solution [50,51]. In this work as it was necessary to use an Hsp27 mutant that does not have a cysteine within the dimer interface of the α-crystallin domain, which further weakens the affinity between subunits [52]. The weaker affinity between Hsp27 subunits compared to αB-c may contribute to the apparent mechanistic differences observed here between Hsp27 and αB-c with regard to their capacity to inhibit the aggregation of FLUC and rhodanese since the increased availability of small, chaperone-active Hsp70 subunits may account for its increased efficacy in inhibiting the aggregation of client proteins compared to αB-c.

### Conclusions

By using a combination of bulk biochemical and single-molecule techniques, we have been able to directly observe, quantify and compare complexes formed by two human sHsps, αB-c and Hsp27, with various model client proteins. The results of this work indicate there are key mechanistic similarities and differences in how these sHsps interact with clients. Both sHsps rapidly form complexes with client proteins that expose significant amounts of hydrophobicity during their aggregation. Our data support a model wherein small sHsp subunits initially bind to client proteins and then additional sHsp subunits are recruited into these complexes over time. In addition, we show that Hsp27 interacts transiently with some clients to inhibit aggregation. Elucidating the mechanisms by which molecular chaperones interact with clients to inhibit aggregation is essential to understanding this highly conserved aspect of proteostasis.

## Materials and methods

### Materials

All common laboratory materials used in this work were purchased from Sigma-Aldrich (St Louis, MO, USA) unless otherwise stated.

### Protein expression and purification

#### Clients

FLUC^ΔCys, K141C, K491C^ (with N-terminal 6x-His tag and C-terminal AviTag; referred to throughout as FLUC) was expressed and purified as previously described [53]. CLIC1^C24^ (an isoform of CLIC1 harbouring mutations of five of the six native cysteines to alanines - C59A, C89A, C178A, C191A, C223A; the remaining cysteine C24 was not modified so it could be used for site-specific fluorescent labelling; this isoform is referred to as CLIC throughout), was expressed and purified as previously described [36]. To express and purify rhodanese, *Escherichia coli* (*E. coli*) BL21 (DE3) cells co-transformed with plasmids encoding biotin ligase (BirA) and SUMO-tagged Avi-rhodanese^K135C, K174C^ (an isoform with cysteine residues replacing the lysines at positions 135 and 174 for site-specific labelling, as well as an N-terminal 6x-His tag and C-terminal AviTag, referred to as rhodanese throughout this work) were used to inoculate a starter culture consisting of LB media supplemented with kanamycin (50 μg/mL) and chloramphenicol (10 μg/mL) antibiotics and grown overnight at 37°C. The starter culture was used to inoculate expression cultures containing LB media supplemented only with kanamycin (50 μg/mL), and the cultures were incubated at 37°C until an OD_600_ of ∼ 0.4 was reached. Expression cultures were then further incubated at 18°C until an OD_600_ of ∼ 0.6-0.8 was reached. To promote the *in vivo* biotinylation of the Avi-tagged recombinant proteins, the media was supplemented with D-biotin (50 μM final concentration) prepared in 10 mM bicine buffer (pH 8.3) prior to protein induction. Expression of recombinant protein was induced by addition of IPTG (0.1 mM) and cultures were incubated on an orbital shaker at 130 rpm overnight (∼ 20 h) at 18°C. The cells were then harvested by centrifugation at 5,000 x *g* for 10 min at 4°C and the pellet stored at −20°C until the recombinant protein was extracted.

Recombinant SUMO-tagged rhodanese was extracted from the bacterial pellet via resuspension in 50 mM Tris-HCl (pH 8.0) supplemented with 300 mM NaCl, 5 mM imidazole, 10% (v/v) glycerol and 20 mM sodium thiosulfate (SUMO IMAC buffer A) that also contained lysozyme (0.5 mg/mL) and EDTA-free cocktail protease inhibitor. The resuspended pellet was then incubated at 4°C for 20 min. The lysates were subjected to probe sonication for 3 min (10 s on/20 s off) at 45% power. The cell debris was then pelleted twice at 24,000 x *g* for 20 min at 4°C and the soluble bacterial lysate passed through a 0.45 μm filter to remove particulates prior to subsequent purification.

The lysate containing SUMO-tagged recombinant rhodanese was first subjected to IMAC chromatography. The bacterial lysate was loaded onto a 5 mL His-Trap Sephadex column pre-equilibrated in SUMO IMAC buffer A. Bound protein was then eluted by addition of the SUMO IMAC buffer A supplemented with 500 mM imidazole (SUMO IMAC buffer B). The fractions containing rhodanese were pooled and dialyzed in the presence of Ulp1 (4 μg/mg of recombinant protein) overnight at 4°C against SUMO IMAC buffer A that did not contain imidazole (SUMO IMAC buffer C). Cleaved recombinant rhodanese was then further purified by passing the dialysed solution over the same His-Trap Sephadex IMAC column equilibrated with SUMO IMAC buffer C and purified as described above, however, this time the recombinant protein did not bind to the column and so was collected in flow through fractions. The presence of recombinant protein in the eluate fractions was confirmed by SDS-PAGE. Fractions containing recombinant protein were pooled and dialyzed into 10 mM Tris (pH 9.0) supplemented with 0.5 mM EDTA and 10% (v/v) glycerol (IEX buffer A) for further purification by ion-exchange (IEX) chromatography. Dialyzed protein (5 mL) was loaded onto a MonoQ 5/50 column pre-equilibrated in IEX buffer A and non-bound proteins eluted from the column with IEX buffer A. Recombinant rhodanese was eluted from the column using a linear salt gradient (0 - 500 mM NaCl) over 20 column volumes. Eluate fractions were analysed by SDS-PAGE for purity, and those fractions containing recombinant rhodanese were pooled and dialyzed into storage buffer (50 mM Tris, pH 7.5, 10 mM MgCl_2_, 5 mM KCl, 10% (v/v) glycerol) overnight at 4°C. Dialyzed protein was concentrated, snap frozen in liquid nitrogen and stored at −80°C until required.

### Molecular chaperones

Plasmids encoding for αB-c_C176_ (i.e. an αB-c isoform with an additional cysteine residue, C176, at the extreme C-terminus that could be used for site-specific labelling of the protein, referred to throughout as αB-c) were transformed into chemically competent *E*. *coli* BL21(DE3) cells and protein expression and purification was performed as previously described [54]. To generate the Hsp27 to be used in this work *E. coli* cells transformed with plasmid encoding for a cysteine mutant of Hsp27 (i.e. Hsp27_C137S,C207_ in which the cysteine in the α-crystallin domain, C137, was replaced with a serine and an additional cysteine was added at the extreme C-terminus, C207, that could be used for site-specific labelling of the protein, referred to throughout as Hsp27) were inoculated in LB containing ampicillin (100 μg/mL, 150 mL) and incubated overnight (37°C) with shaking (200 rpm). This culture was added to expression cultures of LB media containing ampicillin (100 μg/mL), and incubated with shaking until it reached an OD_600_ of 0.6 – 0.8. IPTG was added to a final concentration of 0.25 mM and incubated at 37°C for a further 4 h with constant agitation. Cells were harvested for extraction by centrifugation (5,000 x *g*, 10 min, 4°C). The cell pellet was stored at – 20°C until extraction.

In order to extract recombinant Hsp27, the pellet was resuspended in cell lysis buffer (100 mM Tris–HCl, 10 mM EDTA) containing EDTA-free protease inhibitor cocktail and lysozyme, before incubating on ice for 25 min. The cells were subjected to probe sonication (45% amplitude, 5 s on/10 s off for 3 min) and then centrifuged (24,400 x *g*, 20 min, 4°C). The supernatant was removed and sterile filtered prior to anion-exchange chromatography. The recombinant Hsp27 was purified from contaminating bacterial proteins using anion-exchange and size-exclusion chromatography (SEC). A diethylaminoethyl (DEAE) FF 16/10 anion exchange column (GE Healthcare Lifesciences) equilibrated with DEAE buffer A (20 mM Tris-HCl, pH 8.5, 1 mM EDTA, 0.02% (w/v) NaN_3_) was used as the first step in purification procedures. The cell lysate was loaded onto the column and bound protein eluted using a linear gradient of NaCl from 0 – 200 mM over 8 column volumes by addition of DEAE buffer B (20 mM Tris-HCl, pH 8.5, 1 mM EDTA, 0.02% (w/v) NaN_3_, 2 M NaCl,). The eluate was monitored for protein absorbance at 280 nm and fractions collected from the column analysed by 12% (w/v) SDS-PAGE. Fractions containing Hsp27 were pooled and concentrated prior to loading onto a Sephacryl s300 size-exclusion column (GE Healthcare Lifesciences) previously equilibrated with 50 mM phosphate buffer (pH 7.4, PB). Again, fractions containing the purified Hsp27 were identified by SDS-PAGE, pooled and concentrated. Aliquots were snap frozen in liquid nitrogen prior to storage at −80°C.

### In vitro aggregation assays

The ability of sHsps to prevent the amorphous aggregation of each client protein was assessed by measuring light scatter using *in vitro* aggregation assays. The clients CLIC (30 µM), FLUC (4 µM), and rhodanese (4 µM) were incubated in 50 mM phosphate buffer (pH 7.4) and 10 mM diothiothreitol (DTT) in the absence or presence of varying molar ratios (1:1, 1:2, 1:4, client:sHsp) of αB-c and Hsp27 or the non-chaperone control protein, ovalbumin (1:4, client:ovalbumin). Samples were prepared in duplicate in a clear-bottom 384-well microplate (Greiner Bio-One, Austria) to a final volume of 50 µL per well, and incubated at 42°C without shaking in FLUOstar Optima plate reader (BMG Lab Technologies, Melbourne, Australia) for 20 h (CLIC) or 8 h (FLUC and rhodanese) to cause the amorphous aggregation of the client protein. The light scatter within each sample was continually monitored at 360 nm over the course of the incubation. For each experiment, technical replicates were averaged and used to calculate the relative ability of sHsps to inhibit client aggregation. This was reported as the percentage protection afforded by each sHsp, calculated using equation (1):

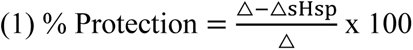

Where △sHsp and △ represent the maximum change in light scatter of the client in the presence and absence (respectively) of sHsp (or control protein) over the course of the incubation. The % protection is reported as the mean ± S.D of 3 independent replicates.

### Bis-ANS assay

Changes in hydrophobicity upon thermal denaturation of client proteins was monitored by 4,4′-Dianilino-1,1′-binaphthyl-5,5′-disulfonic acid (bis-ANS). CLIC, FLUC, or rhodanese (200 nM) were incubated with bis-ANS (20 µM) in 50 mM sodium phosphate buffer (pH 7.4) at room temperature for 10 min. These protein solutions were then dispensed in triplicate into a clear-bottom 384-well microplate (Greiner Bio-One, Austria) to a final volume of 50 µL per well, and incubated at 37°C without shaking in FLUOstar Optima plate reader (BMG Lab Technologies, Melbourne, Australia) for 10 h to promote denaturation of the client, and the bis-ANS fluorescence was measured every 5 min (ex: 355 nm, em: 480 nm). Each emission value was corrected by subtracting a sodium phosphate buffer/bis-ANS blank. The change in bis-ANS fluorescence over time was calculated for each client protein by subtracting the bis-ANS fluorescence value at the first reading, from all corresponding readings after that. The rate of change in bis-ANS fluorescence was calculated by fitting the data to a one-phase association model using GraphPad Prism 9.0 (GraphPad Software Inc.).

### Fluorescent labelling of proteins

Alexa Fluor 488-maleimide (AF488) fluorophore and Alexa Fluor 647-maleimide (AF647) (ThermoFisher Scientific, Waltham, Massachusetts, United States) were used to label the sHsps (Hsp27 and αB-c) and clients (CLIC, FLUC, and Rhodanese), respectively. To do so, 5 mM tris(2-carboxyethyl)phosphine (TCEP) was added to protein (1 mg/mL) to reduce any existing disulphide bonds. Ground ammonium sulfate was dissolved in the protein solution to 40% (w/v) and the sample rotated for 1 h in order to precipitate the protein from solution.

Buffer A (100 mM sodium phosphate, pH 7.4, 200 mM NaCl, 1 mM EDTA, 70% (w/v) ammonium sulfate) and buffer B (100 mM sodium phosphate, pH 7.4, 200 mM NaCl, 1 mM EDTA) were degassed using the Shlenk line technique [55]. Briefly, each buffer was sealed with a rubber plug and a syringe bound to an argon-filled balloon was inserted into the solution. A ‘purge’ syringe was used to degas the buffer for 1 h (buffer A) or 30 min (buffer B).

Following protein precipitation by ammonium sulfate, the solution was centrifuged (20,000 x *g*, 15 min, 4°C) to pellet the precipitated protein and the pellet washed by addition of buffer A (200 μL) followed by centrifugation (20,000 x *g*, 15 min, 4°C). The pellet was then resuspended in buffer B (95 μL) and a five-fold molar excess of fluorophore in DMSO, added such that the final concentration of DMSO was 5% (v/v). The reaction vessel was covered with foil to block light, and mixed via rotation for 2 h at room temperature or, in the case of Hsp27, 37°C overnight.

The sample was loaded onto a Zeba Spin Desalting Column (7K MWCO, 0.5 mL, ThermoFisher Scientific), pre-equilibrated with 50 mM PB (pH 7.4) as per manufacturer instructions. The protein concentration and degree of labelling were determined using the Nanodrop 2000/2000c (ThermoFisher Scientific, Waltham, Massachusetts, United States). Proteins with a degree of labelling that was greater than 90% were used in subsequent experiments.

### Single-molecule TIRF sample preparation

To confirm the presence and composition of complexes formed between clients and sHsps, single-molecule TIRF experiments were performed. AF647-labelled CLIC, FLUC, or rhodanese (2 μM) was incubated with AF488-labelled Hsp27 or αB-c (4 μM) in 50 mM phosphate buffer (pH 7.4) for 7 h at 42°C. Aliquots were taken from the reaction at 0, 0.25, 0.5, 1, 4, and 7 h and crosslinked using bis(sulfosuccinimidyl)suberate (BS^3^) to ensure that the dilution required for single-molecule TIRF imaging did not result in dissociation of any complexes formed.

In order to amine-amine crosslink complexes at each time point, BS^3^ was dissolved in water to 12.5 mM and immediately added to the protein samples to a final concentration of 250 μM, and incubated at room temperature for 30 min. Following incubation, any free crosslinking reagent was quenched via addition of 20 mM Tris hydrochloride (Tris-HCL, pH 7.5) and incubated at room temperature for 15 min before storing at 4°C until imaging.

For imaging, each timepoint sample was diluted to between 1-20 nM into imaging buffer and loaded into flow cells for imaging using TIRF microscopy.

### Calculating the number of fluorophores conjugated per client molecule

To account for the possibility of multiple fluorophores becoming conjugated to the FLUC and rhodanese molecules (due to the two available cysteine residues), the number of fluorophores per labelled FLUC/rhodanese monomer was quantified via single-molecule photobleaching. Rhodanese or FLUC were diluted to 500 pM into 6 mM 6-hydroxy-2,5,7,8-tetramethylchroman-2-carboxylic acid (TROLOX) buffer (imaging buffer) containing oxygen scavenger system (OSS) (a combination of protocatechuic acid (PCA, 2.5 mM) and protocatechuate-3,4-dioxygenase (PCD, 50 nM)) and Guanidine Hydrochloride (2 M) to ensure all proteins were monomeric. A sample (50 μL) was added to a Poly-L-Lysine functionalised coverslip and imaged until all molecules had photobleached. The average number of fluorophores per molecule was calculated as described here for all single-molecule photobleaching analysis. For all subsequent analyses which report the number of subunits within each FLUC/rhodanese molecule (both in complexes and not in complexes), the number of subunits per molecule was divided by the average fluorophore count to correct for any molecules which had multiple fluorophores attached

### Coverslip and flow cell preparation for single-molecule TIRF microscopy

Glass coverslips (24 mm x 24 mm) were cleaned by water bath sonication for 30 min in 100% ethanol followed by 30 min in KOH (5 M). This was repeated once, followed by sonication in Milli-Q water (5 min). Clean coverslips were dried with compressed nitrogen gas.

In order to Poly-L-Lysine functionalise coverslips for quantification of the number of fluorophores conjugated per protein molecule, cleaned coverslips were incubated with Poly-L-Lysine solution (0.01%) (100 μL) in a humidity chamber to prevent drying for 30 min. The coverslips were rinsed with Milli-Q water and dried again with compressed nitrogen gas.

In order to functionalise coverslips for use within flow cells, following the cleaning of coverslips, 5% (v/v) (3-Aminopropyl) triethoxysilane (APTES) (98%, Alfa Aesar, USA) was added to coverslips in Milli-Q water and incubated for 15 min. The coverslips were rinsed with Milli-Q water and dried with compressed nitrogen gas. Methoxyl polyethyleneglycol(mPEG)-succinimidyl valeric acid (SVA) and mPEG-biotin-SVA in 50 mM 3-(Nmorpholino) propanesulfonic acid (MOPS) buffer was added and incubated overnight before rinsing with Milli-Q water and drying again with compressed nitrogen gas. NeutrAvidin (an analog of streptavidin) (ThermoFisher Scientific) was applied directly to the coverslip and incubated for 15 min before rinsing with Milli-Q water and drying with nitrogen gas.

Polydimethylsiloxane (PDMS) custom microfluidics devices were applied directly to the NeutrAvidin functionalised coverslip. Inlets and outlets were inserted at each channel in the PDMS flow cell using PE-20 tubing (Instech, PA, USA) which facilitated the addition of samples into each channel. Polyoxyethylenesorbitanmonolaurate (2% [w/v] Tween20) was incubated within each microfluidic channel and incubated for 30 min to block non-specific protein binding sites and washed out with imaging buffer prior to imaging. For immobilisation of complexes formed with His-tagged CLIC, the flow cell was incubated with a biotinylated anti-6X His tag antibody (cat #ab27025, Abcam, Cambridge, MA) (1 µg/mL) for 10 min and washed with imaging buffer again.

Lastly, crosslinked samples from each time point were diluted into buffer consisting of imaging buffer and an oxygen scavenger system (OSS). The OSS is a combination of protocatechuic acid (PCA, 2.5 mM) and protocatechuate-3,4-dioxygenase (PCD, 50 nM). Samples were incubated within the flow cell for 15 min to bind the coverslip (complexes formed with biotinylated FLUC or rhodanese bound directly to the NeutrAvidin functionalised coverslip), before washing with imaging buffer again prior to image acquisition

### TIRF microscopy instrument setup and data acquisition

All TIRF experiments were performed at room temperature (20°C) using a custom-built system around an inverted microscope (Nikon Eclipse TI). Circularly polarised lasers with constant emission at 488-or 637-nm (200 mW Sapphire, Coherent, Santa Clara, CA, USA) excited samples labelled with AF488 or AF647, respectively. Laser light was directed from a dichroic mirror (ZT405/488/561/635, Semrock, USA) onto the sample through the back-aperture of a 60x 1.49NA TIRF objective (CFI Apochromate TIRF Series 60× objective lens, numerical aperture = 1.49), coated with immersion oil creating an evanescent wave onto coverslip. Emission from the fluorophores passed through the objective, onto the same dichroic mirror. The emission of separate fluorophores was then split using a T635lpxr-UF2 longpass dichroic mirror (Chroma, USA) and then passed through ET525/50m and ET690/50m (Chroma, USA) filters to clean up the emission signal. The final emission signal was then projected onto the electron multiplied charge coupled device (EMCCD) (Andor iXon Life 897, Oxford Instruments, UK) as two separate channels. The camera was operated at −70°C with a pixel distance of 160 nm and electron multiplication gain of 200. Image stacks were recorded with 500 ms exposure time, 600-1000 images per stack, depending on the time taken for 90% of all molecules in the field of view to photobleach completely.

### TIRF data analysis

All TIRF data was first corrected for electronic offset, uneven excitation beam distribution across the field of view, and misalignment of emission channels. Then, using custom written ImageJ scripts [56], each fluorescent molecule was detected, and its’ pixel-wise location within the field of view extracted and compared to the corresponding emission channel for determining colocalisation between the two fluorophores; intensity vs. time trajectories were extracted for each fluorescent spot in order to perform subsequent photobleaching analysis.

Photobleaching analysis was performed using a custom-written analysis package to determine the size of every molecule within each treatment and time point. Briefly, all photobleaching trajectories were classified based on the shape of the data using a residual neural network (ResNet) [57] built using TensorFlow [58] and trained using example photobleaching trajectories extracted from the bleaching of AF647 and AF488 labelled proteins and manually classified prior to training. This model was used to select for trajectories that contained discrete photobleaching steps, and as a quality control step to filter out molecules not suitable for further analysis. Photobleaching step sizes (in fluorescence intensity A.U.) were quantified via change point identification within the trajectories by fitting them to a Bayesian Offline model using the sdt [59] python package. The median size of all final photobleaching steps in all well-defined trajectories were calculated and used in subsequent analysis steps as the size of one photobleaching step (I_step_). The initial intensity of all suitable trajectories was extracted (the average of the first 20 intensity values of each trajectory) and defined as I_initial_. For each molecule, the number of subunits within the fluorescent spot was calculated by I_initial_/I_step_ as described previously [36].

All further filtering and colocalisation analysis was performed using custom-written Python scripts. Plotting used seaborn [60] and matplotlib [61] data visualisation libraries

### Statistics

Statistical analysis of the aggregation assays was performed using GraphPad Prism (9.0). Single-molecule data was analysed using the SciPy [62] or Scikit [63] statistical analysis packages in Python where applicable. Kruskal-Wallis tests with Dunn’s procedure were performed for comparisons between all stoichiometric distributions. A two-way ANOVA with a Tukey’s HSD was performed to compare molecule size data between colocalised and non-colocalised molecules over time. All other specific tests are indicated in the appropriate figure legends. A p-value of less than 0.05 was considered statistically significant, and code used to perform statistical analysis can be found in the code repository.

### Data & Code availability

All scripts used in the analysis workflow described in this work, as well as those that produced the figures and statistical analyses can be found at <u>10.5281/zenodo.10616736.</u>

Example unprocessed data has been provided to explore the analysis workflow described here. Additionally, the data to produce the figures and the statistics has also been provided and both can be found at <u>10.5281/zenodo.10602864.</u>

## Acknowledgements

We would like to thank Prof. Till Böcking (UNSW, Australia) for providing the initial FLUC construct used in this work, and Prof. Benjamin Schuler (University of Zurich, Switzerland) for the initial rhodanese construct. We thank the University of Wollongong staff that provided technical and administrative support for this work. This work was funded by the Australian Research Council (DP220103466). AMvO also acknowledges funding from the National Health and Medical Research Council (APP1197069).

## References

1 Ecroyd, H., Bartelt-Kirbach, B., Ben-Zvi, A., Bonavita, R., Bushman, Y., Casarotto, E., et al. (2023) The beauty and complexity of the small heat shock proteins: a report on the proceedings of the fourth workshop on small heat shock proteins. Cell Stress and Chaperones 28, 621–629 10.1007/s12192-023-01360-x

2 Haslbeck, M., Weinkauf, S. and Buchner, J. (2019) Small heat shock proteins: Simplicity meets complexity. J Biol Chem 294, 2121–2132 10.1074/jbc.REV118.002809

3 Bartelt-Kirbach, B. and Golenhofen, N. (2014) Reaction of small heat-shock proteins to different kinds of cellular stress in cultured rat hippocampal neurons. Cell Stress and Chaperones 19, 145–153 10.1007/s12192-013-0452-9

4 Kato, H., Araki, T., Itoyama, Y., Kogure, K. and Kato, K. (1995) An immunohistochemical study of heat shock protein-27 in the hippocampus in a gerbil model of cerebral ischemia and ischemic tolerance. Neuroscience 68, 65–71 10.1016/0306-4522(95)00141-5

5 Regini, J.W., Ecroyd, H., Meehan, S., Bremmell, K., Clarke, M.J., Lammie, D., et al. (2010) The interaction of unfolding α-lactalbumin and malate dehydrogenase with the molecular chaperone αB-crystallin: a light and X-ray scattering investigation. Mol Vis 16, 2446–2456

6 Stromer, T., Ehrnsperger, M., Gaestel, M. and Buchner, J. (2003) Analysis of the Interaction of Small Heat Shock Proteins with Unfolding Proteins. Journal of Biological Chemistry 278, 18015–18021 10.1074/jbc.M301640200

7 Vendredy, L., Adriaenssens, E. and Timmerman, V. (2020) Small heat shock proteins in neurodegenerative diseases. Cell Stress and Chaperones 25, 679–699 10.1007/s12192-020-01101-4

8. Evgrafov, O.V., Mersiyanova, I., Irobi, J., Van Den Bosch, L., Dierick, I., Leung, C.L., et al. (2004) Mutant small heat-shock protein 27 causes axonal Charcot-Marie-Tooth disease and distal hereditary motor neuropathy. Nat Genet 36, 602–606 10.1038/ng1354

9 Kijima, K., Numakura, C., Goto, T., Takahashi, T., Otagiri, T., Umetsu, K., et al. (2005) Small heat shock protein 27 mutation in a Japanese patient with distal hereditary motor neuropathy. J Hum Genet 50, 473–476 10.1007/s10038-005-0280-6

10 Datskevich, P.N., Nefedova, V.V., Sudnitsyna, M.V. and Gusev, N.B. (2012) Mutations of small heat shock proteins and human congenital diseases. Biochemistry Moscow 77, 1500–1514 10.1134/S0006297912130081

11 Capponi, S., Geuens, T., Geroldi, A., Origone, P., Verdiani, S., Cichero, E., et al. (2016) Molecular chaperones in the pathogenesis of Amyotrophic Lateral Sclerosis: the role of HSPB1. Human Mutation 37, 1202–1208 10.1002/humu.23062

12 Tóth, M.E., Szegedi, V., Varga, E., Juhász, G., Horváth, J., Borbély, E., et al. (2013) Overexpression of Hsp27 ameliorates symptoms of Alzheimer’s disease in APP/PS1 mice. Cell Stress and Chaperones 18, 759–771 10.1007/s12192-013-0428-9

13 Litt, M. (1998) Autosomal dominant congenital cataract associated with a missense mutation in the human alpha crystallin gene CRYAA. Human Molecular Genetics 7, 471– 474 10.1093/hmg/7.3.471

14 Raju, I. and Abraham, E.C. (2013) Mutants of human αB-crystallin cause enhanced protein aggregation and apoptosis in mammalian cells: Influence of co-expression of HspB1. Biochemical and Biophysical Research Communications 430, 107–112 10.1016/j.bbrc.2012.11.051

15 Xia, X.-Y., Wu, Q.-Y., An, L.-M., Li, W.-W., Li, N., Li, T.-F., et al. (2014) A novel P20R mutation in the alpha-B crystallin gene causes autosomal dominant congenital posterior polar cataracts in a Chinese family. BMC Ophthalmol 14, 108 10.1186/1471-2415-14-108

16 Liu, Y., Zhang, X., Luo, L., Wu, M., Zeng, R., Cheng, G., et al. (2006) A novel αB-Crystallin mutation associated with autosomal dominant congenital lamellar cataract. Investigative Ophthalmology & Visual Science 47, 1069–1075 10.1167/iovs.05-1004

17 Delbecq, S.P. and Klevit, R.E. (2019) HSPB5 engages multiple states of a destabilized client to enhance chaperone activity in a stress-dependent manner. J Biol Chem 294, 3261–3270 10.1074/jbc.RA118.003156

18 Braun, N., Zacharias, M., Peschek, J., Kastenmüller, A., Zou, J., Hanzlik, M., et al. (2011) Multiple molecular architectures of the eye lens chaperone αB-crystallin elucidated by a triple hybrid approach. Proc Natl Acad Sci U S A 108, 20491–20496 10.1073/pnas.1111014108

19 Peschek, J., Braun, N., Franzmann, T.M., Georgalis, Y., Haslbeck, M., Weinkauf, S., et al. (2009) The eye lens chaperone α-crystallin forms defined globular assemblies. Proc Natl Acad Sci U S A 106, 13272–13277 10.1073/pnas.0902651106

20 Jovcevski, B., Kelly, M.A., Rote, A.P., Berg, T., Gastall, H.Y., Benesch, J.L.P., et al. (2015) Phosphomimics destabilize Hsp27 oligomeric assemblies and enhance chaperone activity. Chemistry & Biology 22, 186–195 10.1016/j.chembiol.2015.01.001

21 Aquilina, J.A., Shrestha, S., Morris, A.M. and Ecroyd, H. (2013) Structural and functional aspects of hetero-oligomers formed by the small heat shock proteins αB-Crystallin and Hsp27 *. Journal of Biological Chemistry 288, 13602–13609 10.1074/jbc.M112.443812

22 Sha, E., Nakamura, M., Ankai, K., Yamamoto, Y.Y., Oka, T. and Yohda, M. (2019) Functional and structural characterization of HspB1/Hsp27 from Chinese hamster ovary cells. FEBS Open Bio 9, 1826–1834 10.1002/2211-5463.12726

23. Feil, I.K., Malfois, M., Hendle, J., Zandt, H. van der and Svergun, D.I. (2001) A Novel quaternary structure of the dimeric α-Crystallin domain with chaperone-like activity *. Journal of Biological Chemistry 276, 12024–12029 10.1074/jbc.M010856200

24 Aquilina, J.A., Benesch, J.L.P., Ding, L.L., Yaron, O., Horwitz, J. and Robinson, C.V. (2005) Subunit exchange of polydisperse proteins: mass spectrometry reveals consequences of αA-crystallin truncation *. Journal of Biological Chemistry 280, 14485–14491 10.1074/jbc.M500135200

25 Hayes, D., Napoli, V., Mazurkie, A., Stafford, W.F. and Graceffa, P. (2009) Phosphorylation dependence of Hsp27 multimeric size and molecular chaperone function *. Journal of Biological Chemistry 284, 18801–18807 10.1074/jbc.M109.011353

26 Stengel, F., Baldwin, A.J., Bush, M.F., Hilton, G.R., Lioe, H., Basha, E., et al. (2012) Dissecting heterogeneous molecular chaperone complexes using a mass spectrum deconvolution approach. Chemistry & Biology 19, 599–607 10.1016/j.chembiol.2012.04.007

27 Eyles, S.J. and Gierasch, L.M. (2010) Nature’s molecular sponges: Small heat shock proteins grow into their chaperone roles. Proc Natl Acad Sci U S A 107, 2727–2728 10.1073/pnas.0915160107

28 Clouser, A.F. and Klevit, R.E. (2017) pH-dependent structural modulation is conserved in the human small heat shock protein HSBP1. Cell Stress Chaperones 22, 569– 575 10.1007/s12192-017-0783-z

29 Liu, Z., Wang, C., Li, Y., Zhao, C., Li, T., Li, D., et al. (2018) Mechanistic insights into the switch of αB-crystallin chaperone activity and self-multimerization. J Biol Chem 293, 14880–14890 10.1074/jbc.RA118.004034

30 Santhanagopalan, I., Degiacomi, M.T., Shepherd, D.A., Hochberg, G.K.A., Benesch, J.L.P. and Vierling, E. (2018) It takes a dimer to tango: Oligomeric small heat shock proteins dissociate to capture substrate. J Biol Chem 293, 19511–19521 10.1074/jbc.RA118.005421

31 Haslbeck, M., Walke, S., Stromer, T., Ehrnsperger, M., White, H.E., Chen, S., et al. (1999) Hsp26: a temperature-regulated chaperone. EMBO J 18, 6744–6751 10.1093/emboj/18.23.6744

32 Mymrikov, E.V., Daake, M., Richter, B., Haslbeck, M. and Buchner, J. (2017) The chaperone activity and substrate spectrum of human small heat shock proteins. J Biol Chem 292, 672–684 10.1074/jbc.M116.760413

33 Lee, G.J., Roseman, A.M., Saibil, H.R. and Vierling, E. (1997) A small heat shock protein stably binds heat-denatured model substrates and can maintain a substrate in a folding-competent state. EMBO J 16, 659–671 10.1093/emboj/16.3.659

34. Johnston, C.L., Marzano, N.R., Van Oijen, A.M. and Ecroyd, H. (2018) Using single-molecule approaches to understand the molecular mechanisms of heat-shock protein chaperone function. Journal of Molecular Biology 430, 4525–4546 10.1016/j.jmb.2018.05.021

35. Rice, L.J., Ecroyd, H. and Van Oijen, A.M. (2021) Illuminating amyloid fibrils: fluorescence-based single-molecule approaches. Computational and Structural Biotechnology Journal 19, 4711–4724 10.1016/j.csbj.2021.08.017

36. Johnston, C.L., Marzano, N.R., Paudel, B.P., Wright, G., Benesch, J.L.P., van Oijen, A.M., et al. (2020) Single-molecule fluorescence-based approach reveals novel mechanistic insights into human small heat shock protein chaperone function. J Biol Chem 296, 100161 10.1074/jbc.RA120.015419

37 Haslbeck, M. (1999) Hsp26: a temperature-regulated chaperone. The EMBO Journal 18, 6744–6751 10.1093/emboj/18.23.6744

38 lindner, r.a., kapur, a. and carver, j.a. (1997) The interaction of the molecular chaperone, α-Crystallin, with molten globule states of bovine α-lactalbumin. J Biol Chem 272, 27722–27729 10.1074/jbc.272.44.27722

39 Inoue, R., Takata, T., Fujii, N., Ishii, K., Uchiyama, S., Sato, N., et al. (2016) New insight into the dynamical system of αB-crystallin oligomers. Sci Rep 6, 29208 10.1038/srep29208

40 Finn, T.E., Nunez, A.C., Sunde, M. and Easterbrook-Smith, S.B. (2012) Serum albumin prevents protein aggregation and amyloid formation and retains chaperone-like activity in the presence of physiological ligands. J Biol Chem 287, 21530–21540 10.1074/jbc.M112.372961

41 Marini, I., Moschini, R., Corso, A.D. and Mura, U. (2005) Chaperone-like features of bovine serum albumin: a comparison with α-crystallin. Cell. Mol. Life Sci. 62, 3092–3099 10.1007/s00018-005-5397-4

42 Delbecq, S.P. and Klevit, R.E. (2013) One size doesn’t fit all: the oligomeric states of αB Crystallin. FEBS Lett 587, 10.1016/j.febslet.2013.01.021

43 Tiroli-Cepeda, A.O. and Ramos, C.H.I. (2010) Heat causes oligomeric disassembly and increases the chaperone activity of small heat shock proteins from sugarcane. Plant Physiology and Biochemistry 48, 108–116 10.1016/j.plaphy.2010.01.001

44 Scheidt, T., Carozza, J.A., Kolbe, C.C., Aprile, F.A., Tkachenko, O., Bellaiche, M.M.J., et al. (2021) The binding of the small heat-shock protein αB-crystallin to fibrils of α-synuclein is driven by entropic forces. Proc. Natl. Acad. Sci. U.S.A. 118, e2108790118 10.1073/pnas.2108790118

45 Weids, A.J., Ibstedt, S., Tamás, M.J. and Grant, C.M. (2016) Distinct stress conditions result in aggregation of proteins with similar properties. Sci Rep 6, 24554 10.1038/srep24554

46 Yu, C., Leung, S.K.P., Zhang, W., Lai, L.T.F., Chan, Y.K., Wong, M.C., et al. (2021) Structural basis of substrate recognition and thermal protection by a small heat shock protein. Nat Commun 12, 3007 10.1038/s41467-021-23338-y

47 Miller, A.P., O’Neill, S.E., Lampi, K.J. and Reichow, S.L. (2023) The α-Crystallin chaperones undergo a quasi-ordered co-aggregation process in response to saturating client interaction. Biophysics; 2023 10.1101/2023.08.15.553435

48 Gonçalves, C.C., Sharon, I., Schmeing, T.M., Ramos, C.H.I. and Young, J.C. (2021) The chaperone HSPB1 prepares protein aggregates for resolubilization by HSP70. Sci Rep 11, 17139 10.1038/s41598-021-96518-x

49 Benesch, J.L.P., Aquilina, J.A., Baldwin, A.J., Rekas, A., Stengel, F., Lindner, R.A., et al. (2010) The quaternary organization and dynamics of the molecular chaperone Hsp26 are thermally regulated. Chemistry & Biology 17, 1008–1017 10.1016/j.chembiol.2010.06.016

50 Skouri-Panet, F., Michiel, M., Férard, C., Duprat, E. and Finet, S. (2012) Structural and functional specificity of small heat shock protein HspB1 and HspB4, two cellular partners of HspB5: Role of the in vitro hetero-complex formation in chaperone activity. Biochimie 94, 975–984 10.1016/j.biochi.2011.12.018

51 Bova, M.P., Mchaourab, H.S., Han, Y. and Fung, B.K.-K. (2000) Subunit exchange of small heat shock proteins. J Biol Chem 275, 1035–1042 10.1074/jbc.275.2.1035

52 Alderson, T.R., Roche, J., Gastall, H.Y., Dias, D.M., Pritišanac, I., Ying, J., et al. (2019) Local unfolding of the HSP27 monomer regulates chaperone activity. Nat Commun 10, 1068 10.1038/s41467-019-08557-8

53. Marzano, N.R., Paudel, B.P., Van Oijen, A.M. and Ecroyd, H. (2022) Real-time single-molecule observation of chaperone-assisted protein folding. Sci. Adv. 8, eadd0922 10.1126/sciadv.add0922

54 Horwitz, J., Huang, Q.-L., Ding, L. and Bova, M.P. (1998) Lens α-crystallin: Chaperone-like properties. Methods in Enzymology. Elsevier; 1998. p. 365–383. 10.1016/S0076-6879(98)90032-5

55 Chandra, T. and Zebrowski, J.P. (2014) Reactivity control using a Schlenk line. *J*. Chem. Health Saf. 21, 22–28 10.1016/j.jchas.2014.02.001

56 Schneider, C.A., Rasband, W.S. and Eliceiri, K.W. (2012) NIH Image to ImageJ: 25 years of image analysis. Nat Methods 9, 671–675 10.1038/nmeth.2089

57 He, K., Zhang, X., Ren, S. and Sun, J. (2015) Deep residual learning for image recognition 10.48550/ARXIV.1512.03385

58 TensorFlow Developers (2023) TensorFlow 10.5281/ZENODO.4724125

59 Lschr and Luhk (2023) schuetzgroup/sdt-python: v18.1 10.5281/ZENODO.8239822

60 Waskom, M. (2021) seaborn: statistical data visualization. JOSS 6, 3021 10.21105/joss.03021

61 Thomas A Caswell, Elliott Sales de Andrade, Antony Lee, Michael Droettboom, Tim Hoffmann, Jody Klymak, et al. (2023) matplotlib/matplotlib: REL: v3.7.4 10.5281/ZENODO.10152802

62 Virtanen, P., Gommers, R., Oliphant, T.E., Haberland, M., Reddy, T., Cournapeau, D., et al. (2020) SciPy 1.0: fundamental algorithms for scientific computing in Python. Nat Methods 17, 261–272 10.1038/s41592-019-0686-2

63 Terpilowski, M. (2019) scikit-posthocs: Pairwise multiple comparison tests in Python. JOSS 4, 1169 10.21105/joss.01169

