## Supplementary Figures 1-4 for "Single-molecule observations of human small heat shock proteins in complex with aggregation-prone client proteins"

### Supplementary Material

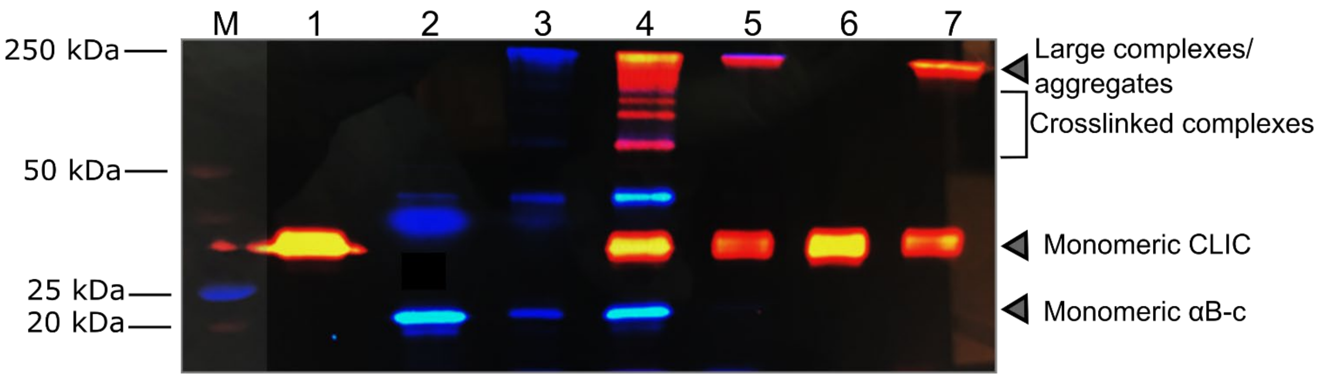


**Supplementary figure 1. Chemical crosslinking maintains sHsp oligomer size and prevents sHsp:client dissociation**. Samples containing AF488-labelled αB-c-(2 µM) or AF647-labelled CLIC (1 µM) were incubated in the absence and presence of one another (4 h, 42°C). Samples were chemically crosslinked using 20 x molar excess of $BS^3$. Samples were subjected to SDS-PAGE and imaged for the corresponding fluorophore to confirm crosslinking, and compared to non-crosslinked controls. Gel shows non-crosslinked CLIC (1), non-crosslinked αB-c (2), crosslinked αB-c (3), αB-c and CLIC incubated together and crosslinked (4), crosslinked αB-c, incubated with CLIC and then crosslinked following incubation (5), crosslinked, unheated CLIC (6), incubated and crosslinked CLIC (7).

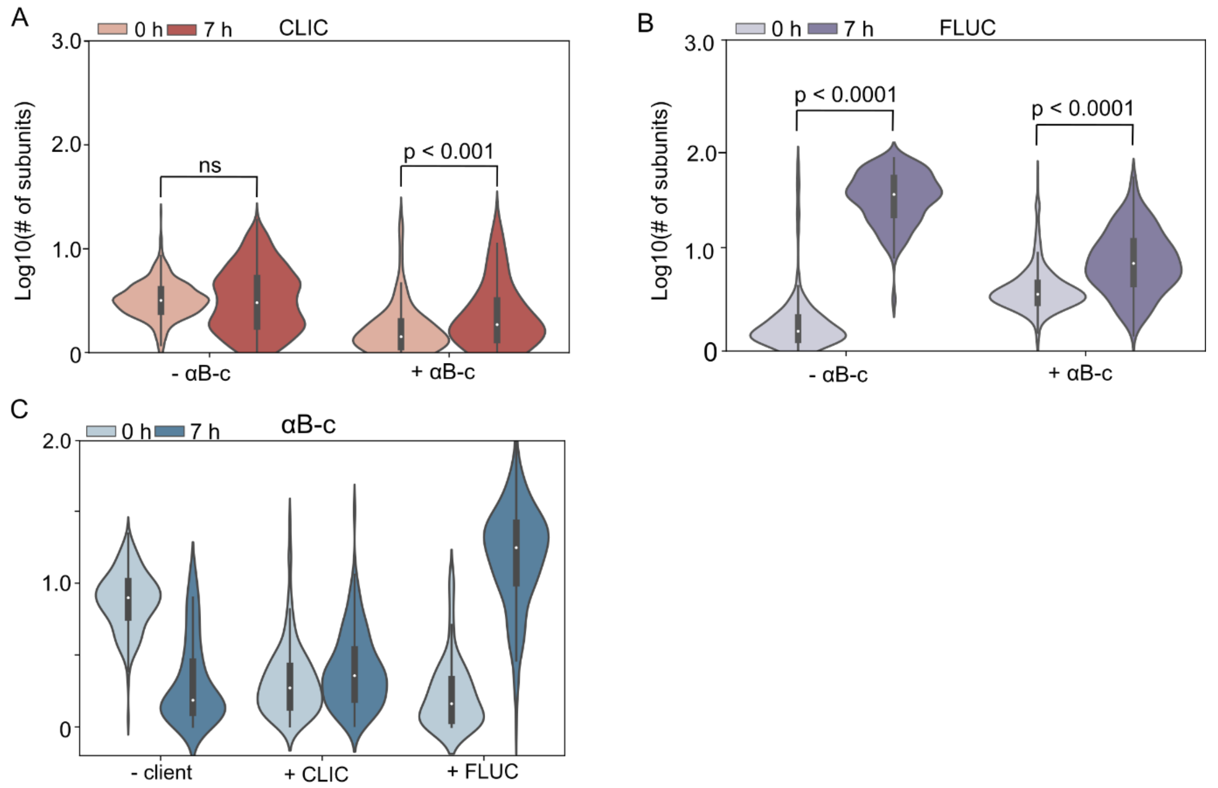


**Supplementary figure 2. αB-c inhibits aggregation of CLIC- or FLUC- AF647 and the increase in size of αB-c molecules is client protein-dependent**. CLIC- or FLUC-AF647 were incubated (42°C for up to 7 h) in the absence or presence αB-c (2:1 molar ratio); aliquots were taken at the start (0 h) and end (7 h) of the incubation. Following incubation, samples were crosslinked, diluted, and incubated in flow cells for 15 min before imaging using TIRF microscopy. (**A-C**) Violin plots show the distributions of all (colocalised and non-colocalised) molecule sizes ($\log_{10}$ number of subunits/molecule) prior to and after incubation of (A) CLIC-AF647 and (B) FLUC-AF647 with ('+αB-c') or without ('-αB-c') αB-c-AF488; or, (C) αB-c-AF488 alone ('-client') and in the presence of CLIC–AF647('+ CLIC') or FLUC-AF647 ('FLUC'). Results include measurements from 3 independent experiments and, where marked, statistical comparison between distributions was performed via Kruskal-Wallis test for multiple comparisons with Dunn's procedure (p values indicated).

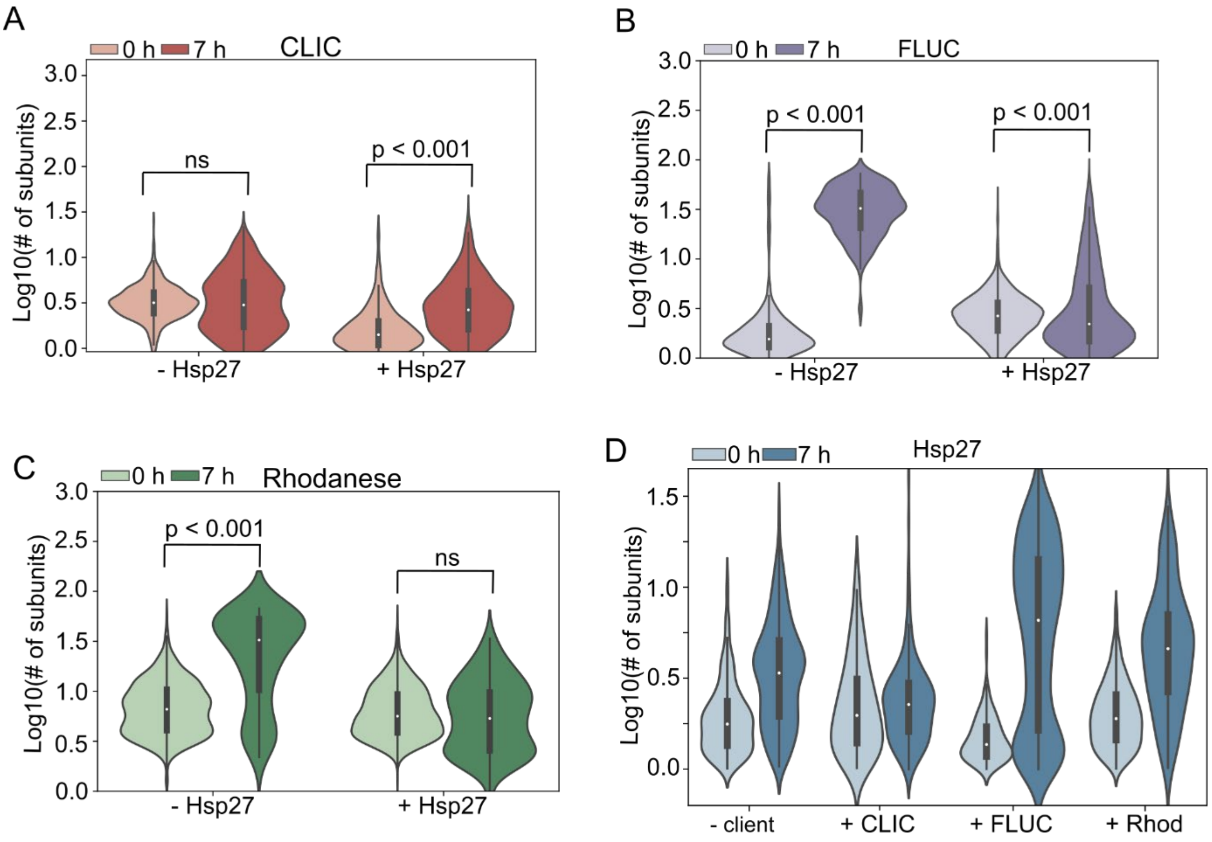


**Supplementary figure 3. Hsp27 inhibits aggregation of CLIC-, FLUC- and rhodanese-AF647, and Hsp27 molecule size is client-protein dependent**. CLIC-, FLUC- and rhodanese-AF647 were incubated (42°C for 7 h) in the presence (2:1 molar ratio) and absence of one another; aliquots were taken at the start (0 h) and end (7 h) of the incubation. Following incubation, samples were crosslinked, diluted, and incubated in flow cells for 15 min before imaging using TIRF microscopy. (**A-D**) Violin plots show the distributions of all (colocalised and non-colocalised) molecule sizes ($\log_{10}$ number of subunits/molecule) prior to and after incubation of (A) CLIC-AF647, (B) FLUC-AF647, and (C) rhodanese-AF647 with ('+sHsp') or without ('-sHsp') Hsp27-AF488; or, (C) Hsp27-AF488 alone ('-client') and in the presence of CLIC-AF647 ('+ CLIC'), FLUC-AF647 ('FLUC'), or rhodanese-AF647 ('Rhodanese'). Results include measurements from 3 independent experiments and, where marked, statistical comparison between distributions was performed via Kruskal-Wallis test for multiple comparisons with Dunn's procedure.

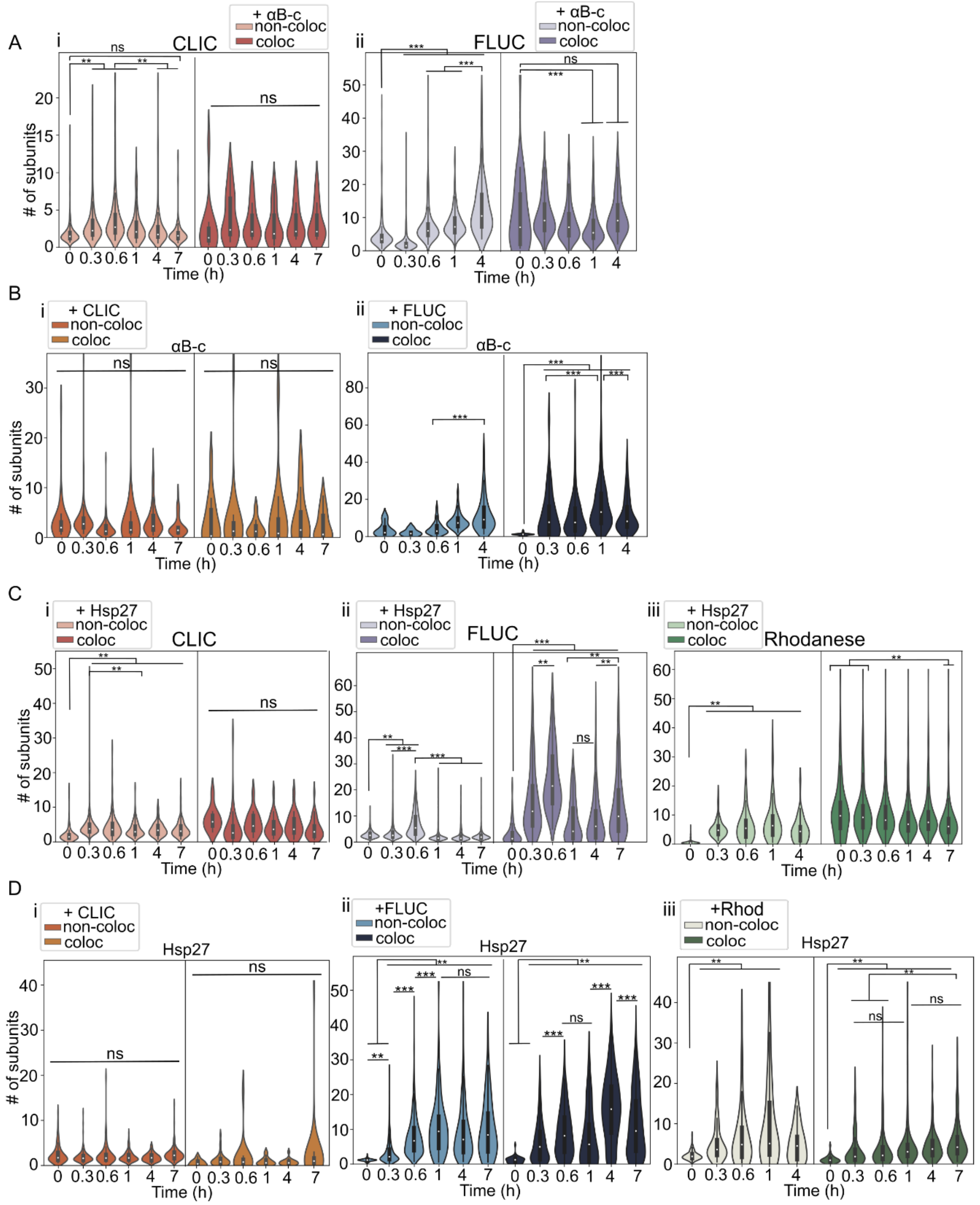


**Supplementary Figure 4. sHsps maintain client proteins in small oligomeric states.** AF488-labelled sHsps (αB-c or Hsp27) were incubated with the AF647-labelled client proteins (CLIC, FLUC or rhodanese) (42°C for 7 h, 2:1 molar ratio) and aliquots were taken throughout the incubation. Samples were immediately crosslinked, diluted, and incubated in flow cells for 15 min before imaging using TIRF microscopy. The molecule size (# of subunits/molecule) was calculated for all molecules, and the data was filtered for the **(A, C)** client and **(B, D)** sHsp molecules that were colocalised with one another (i.e., in complexes) and those that were not colocalised (i.e., not in complexes). (A,C) Violin plots showing the distribution of the number of subunits per molecule of both colocalised ('Coloc') and non-colocalised ('Non-coloc') molecules at each time point for the clients CLIC (i, red), FLUC (ii, purple) and/or rhodanese (iii, green) incubated with (B) αB-c or (D) Hsp27. The size of the sHsps in these treatments are shown in the corresponding plots for (B) αB-c and (D) Hsp27. The data shown contains the number of subunits from all molecules in 3 independent experiments. A two-way ANOVA was performed for all treatments, and, where relevant, statistical differences are marked (** = $p < 0.005$, *** = $p < 0.0005$, ns indicates no significant difference. *The median and error values presented in Figure 4 (main text) are calculated from all molecules depicted here as distributions.*
